# Modeling and Design of Chitosan-PCL Bi-Layered Microspheres for Intravitreal Controlled Release

**DOI:** 10.1101/2024.01.11.575289

**Authors:** Eduardo A. Chacin Ruiz, Samantha L. Carpenter, Katelyn E. Swindle-Reilly, Ashlee N. Ford Versypt

## Abstract

Chronic retinal diseases usually require repetitive local dosing. Depending on factors such as dosing frequency, mode of administration, and associated costs, this can result in poor patient compliance. A better alternative involves using controlled release drug delivery systems to reduce the frequency of intravitreal dosing and extend drug release. However, reaching the market stage is a time-consuming process. In this study, we employed two computational approaches to model and estimate the parameters governing the diffusion-controlled drug release of bovine serum albumin and bevacizumab (an agent that slows neovascularization due to retinal disorders) from bi-layered core-shell microspheres composed of chitosan and polycaprolactone (PCL). We used the estimated parameters to simulate the cumulative release under various conditions, optimize device design to guide future experimental efforts and improve the duration of release above a target daily therapeutic release rate from the microspheres. We investigated the effects of polymeric layer sizes on drug release. We provided straightforward computational tools for others to reuse in designing bi-layered microspheres suitable for addressing intravitreal drug delivery needs in the treatment of ocular neovascularization in chronic retinal diseases.

## 1. Introduction

Chronic diseases usually require repeated dosing to maintain therapeutic drug concentrations over extended periods to manage symptoms and prevent disease progression. This is particularly challenging in the treatment of chronic retinal disorders like age-related macular degeneration (AMD) and diabetic retinopathy, which involve neovascularization and require repetitive intravitreal injections. These frequent injections can cause fear and anxiety in patients and are associated with an increased risk of ocular complications [1]. Therefore, there is a critical need for drug delivery systems (DDSs) capable of maintaining therapeutic concentrations for extended periods, thereby improving patient compliance compared to classic formulations for administering drugs directly through systemic or localized routes.

Achieving long-term, sustained delivery at therapeutic levels using DDSs requires consideration of the parameters that govern the drug release dynamics. In layered polymeric DDSs, the parameters include drug diffusion coefficients, burst release percentage, and drug partition coefficients, which are not necessarily known beforehand; however, they influence the amount and rate of delivery at the target locations. Experimental testing of every possible design combination is impractical due to time and cost constraints. Mathematical modeling of drug release from DDSs enables predictive simulations of drug release behavior across a vast design space. These models can estimate the key parameters and optimize device design, reducing the amount of experimentation required while improving the treatment of chronic diseases.

Several mechanisms can govern drug release from polymeric DDSs, including diffusion, swelling, and erosion [2–5]. Here, we focus on diffusion-controlled release where diffusion through the polymer(s) dominates the drug transport. Diffusion-controlled release is modeled by Fick’s second law, often assuming a constant diffusion coefficient in each material, perfect sink conditions in the medium, and negligible swelling and erosion of the DDS [4, 5]. Analytical solutions exist for single-layered DDSs with ideal geometries [6–11] and multi-layered spherical DDSs under certain idealized conditions [12–15], which are reviewed in Chacin Ruiz et al. [2]. Unfortunately, the analytical solutions are complicated to evaluate as they typically involve infinite sums of trigonometric functions. While these solutions rely on fixed values or assumed ratios among parameters, fewer studies have focused on parameter estimation from such single-layered [16–18] or multi-layered spheres [19, 20]. For evaluating cases that can be solved by analytical techniques, extending to other conditions without analytical solutions, connecting with parameter estimation software tools, and considering DDSs with more elaborate geometries, we turned to numerical approaches. We used a finite difference approach in MATLAB for symmetric multi-layered spheres. We also used an alternative finite element approach in COMSOL to enable future extensions to layered geometries of other shapes and loading and boundary conditions that are not easily modeled in MATLAB.

This work addresses the challenge of improving the design of intravitreal DDSs through computational modeling, thereby reducing the design space for costly and time-consuming experiments. We present numerical approaches based on finite differences and finite elements for solving the long-term drug release problem and estimating important parameters in diffusion-controlled core-shell spherical DDSs. As a case study, we applied these numerical methods to chitosan-polycaprolactone (PCL) microspheres releasing bovine serum albumin (BSA) and bevacizumab [21], an agent that slows neovascularization in cancer and eye disorders. The calculated parameters were estimated from published experimental data and used to explore the effects of the design variables–the sizes of the polymer layers–on drug release metrics of cumulative release percentage and daily drug release rate. This framework supports the rational design of intravitreal DDSs to meet specific therapeutic thresholds, providing a generalizable toolset for both experimentalists and theoreticians working on sustained DDSs.

## 2. Methods

### 2.1. Geometry

The DDS used here is a core-shell microsphere (Figure 1). A single dose includes many of the microspheres, which are assumed to behave identically. The radius of the core is *R*_*core*_, and that of the shell is *R*_*shell*_. The shell thickness is denoted Δ*R* = *R*_*shell*_ − *R*_*core*_. The minΔ*R* = 0, thus *R*_*shell*_ ≥ *R*_*core*_. Note that Δ*R* = 0 is just the single-layered microsphere case, where *R*_*shell*_ = *R*_*core*_. Radial symmetry about the center of each microsphere is assumed.

**Figure 1:**
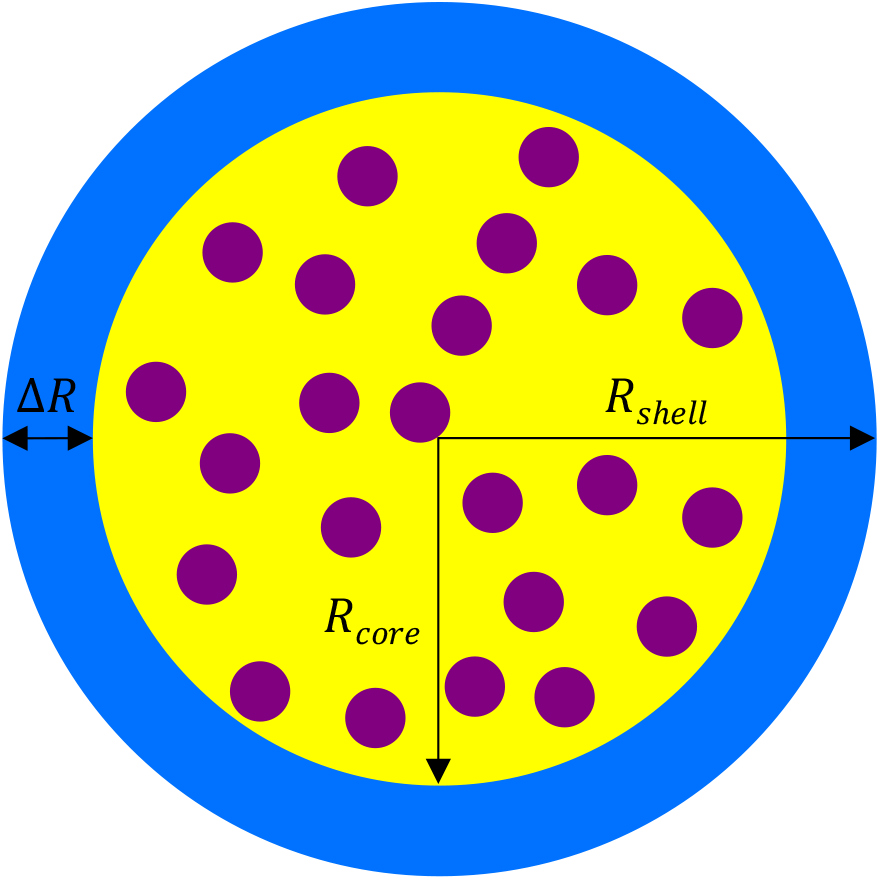
2D cross-section of a core-shell microsphere. A drug is loaded only in the core. Yellow: core. Blue: shell. Purple: drug. *R*: radius. Δ*R* = *R*_*shell*_ *−R*_*core*_. Note that the drug is not illustrated to scale.

As test cases for demonstrating our computational tools for designing DDSs for intravitreal controlled release, we used data from an experimental study from our team [21], in which the DDS consisted of a drug-loaded core made of chitosan and a PCL outer shell to protect the chitosan core from degradation and extend drug release. PCL has a very slow hydrolytic degradation rate [22– 24]. Bartnikowski et al. [24] combined data from multiple studies in the literature showing that less than 5% of PCL mass was lost by 10 months of *in vitro* degradation in cell-free media at body temperature; Lam et al. [25] showed that the 5% loss threshold was not passed until nearly 2 years. This is much longer than the 6-month time scale in the present study and in Jiang et al. [21]; thus, PCL is assumed not to degrade. Chitosan was shown to degrade by 6 months when not surrounded by a PCL outer shell, but it was protected when a shell was present [21]. Thus, we do not consider the case of Δ*R* = 0 (chitosan single-layered microsphere) as a test case for the diffusion-controlled model. Swelling is also assumed to be negligible for both polymers. Thus, diffusion-controlled release is the dominant release mechanism. In Jiang et al. [21], the mean *R*_*core*_ = 5.10 µm of chitosan, and the mean Δ*R* = 1.25 µm of PCL. Two drugs were used in Jiang et al. [21] and are considered here: BSA and bevacizumab.

### 2.2 Mathematical model

Fick’s second law for time-dependent diffusion is

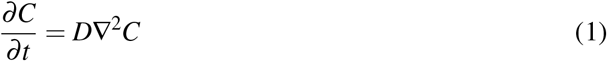

where 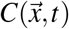 is the drug concentration, 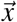 refers to the spatial coordinates in either Cartesian or spherical coordinate systems, *t* is time, and *D* is a uniform diffusion coefficient. For two concentric layers of a symmetric sphere, this becomes

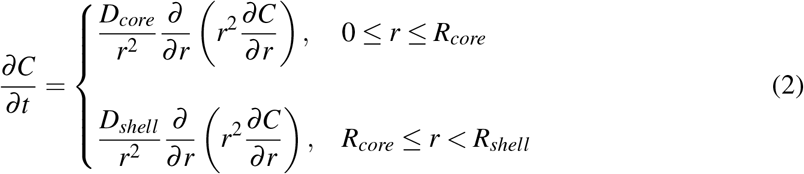

where *D*_*core*_ and *D*_*shell*_ are the diffusion coefficients for the core and shell layers, respectively, and *r* is the position along the radial dimension.

For the bi-layered or core-shell sphere, there are three boundary conditions to define for *t >* 0 at *r* = 0, *R*_*core*_, and *R*_*shell*_. At the center of the sphere (*r* = 0), the radial symmetry boundary condition is applied:

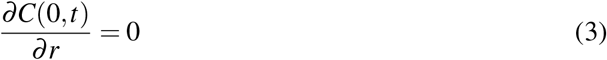

The condition at the interface (*r* = *R*_*core*_) between the two different polymeric domains is the flux continuity condition while allowing for drug partitioning preferentially between the layers based on material properties:

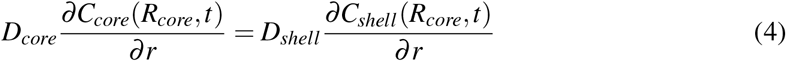

subject to

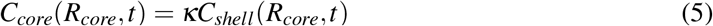

where *C*_*core*_(*R*_*core*_,*t*) is the drug concentration in the core at the interface, *κ* is the partition coefficient, and *C*_*shell*_(*R*_*core*_,*t*) is the drug concentration in the shell at the interface. At the external surface of the sphere (*r* = *R*_*shell*_), the perfect sink condition is assumed: *C*(*R*_*shell*_,*t*) = 0. This means that all the drug released is immediately cleared away, which is a common assumption in biological tissues.

The initial concentration for the drug-loaded core and non-loaded shell is

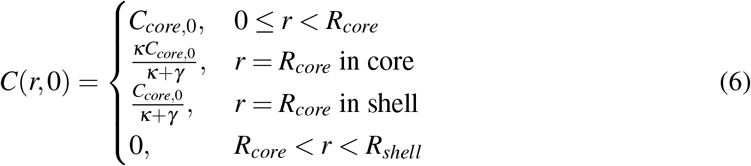

where *C*_*core*,0_ is the initial concentration inside the core, which is set as *C*_*core*,0_ = 1 arbitrary units (a.u.) for normalizing the concentration calculations. Note this normalized value is the concentration that remains immediately after the burst release, which we assumed to be instantaneous at *t* = 0. The initial values at the interface are derived in Section S1 in the Supplementary Material as the weighted averages of the initial values in the core and the shell, with the weighting dependent on the partition coefficient *κ* and the transport properties in the layers (Equations (S26) and (S27) in the Supplementary Material). *γ* is a ratio of the diffusion coefficients and the grid spacing in the two layers and is defined in Equation (S18) in the Supplementary Material.

Drug release is quantified in terms of cumulative release, as this metric is often reported in experimental studies. The cumulative release is calculated by

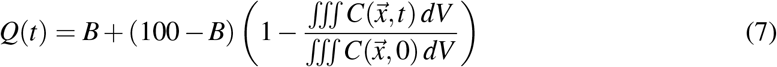

where *Q*(*t*) is the cumulative percentage of drug released as a function of time, *B* is the burst release or the percentage of the total drug loaded that is released immediately upon *in vivo* administration or placement into an *in vitro* medium, *V* is volume, and the last term refers to the fractional amount of drug remaining in the DDS at each time relative to that present in the DDS initially after the burst release. The total amount of drug in the core-shell microsphere at any time is calculated as the volume-integral of the spatiotemporal distribution of 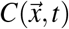 from the solution of Equations (2)– (6).

The amount of drug released over a time interval *A*_*rel*_(*t*) is determined using the cumulative release from each microsphere as

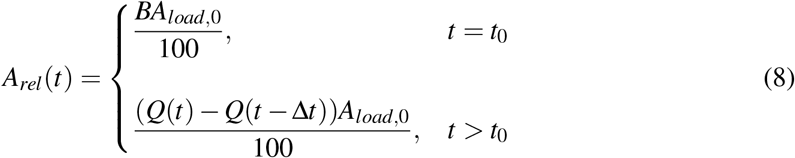

where *t*_0_ denotes the initial time, Δ*t* is the time interval, and *A*_*load*,0_ corresponds to the initial amount (in moles) of drug loaded into a dose of multiple microspheres. We assumed that each microsphere has a uniform *C*_*core*,0_ in its core. *C*_*core*,0_ for diffusion and the cumulative release calculations is dimensionless and set to 1 to normalize by the dimensional initial concentration. Additionally, we assumed that the dimensional initial concentration in each microsphere core in units of moles per volume is the same across all microspheres as this represents the loading capacity that is a material property. Microspheres of different core radii have different volumes over which the drug is distributed. Thus, we assumed that a sufficient number of microspheres comprise a dose to reach the *A*_*load*,0_ amount. Different drugs have different molecular weights, and these were used when converting from initial amounts in mass units to mole units.

The release rate *Ȧ*_*rel*_ (*t*) can be expressed as

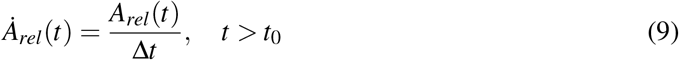

which gives the release rate between two time points spaced by Δ*t*, or the approximation to the derivative of the cumulative release curve. The release rate is fastest at early times. As the Δ*t* needed for burst release at time *t* = 0 is assumed infinitesimally short, the instantaneous burst release rate is theoretically infinitely high (practically, it must be finite). Thus, Equation (9) is undefined at *t* = 0, and we only considered the finite drug release rates calculated for *t* > 0.

### 2.3. Numerical methods

We solved the model for drug release from the core-shell microsphere DDS defined in Equations (2)–(6) numerically using two alternative approaches: finite differences in MATLAB and finite elements in COMSOL Multiphysics.

#### 2.3.1. Finite difference approach in MATLAB

For the radially symmetric core-shell sphere described in Section 2.1 with uniform diffusion coefficients in each layer and the initial and boundary conditions described in Section 2.2, the finite difference approach is relatively straightforward to implement in technical computing languages such as MATLAB or Python. Here, we used MATLAB. The first step in the approach is to normalize the spherical domain by scaling by *R*_*shell*_ so that the domain is 0 ≤ *r/R*_*shell*_ ≤ 1. Then, a variable *α* is defined as

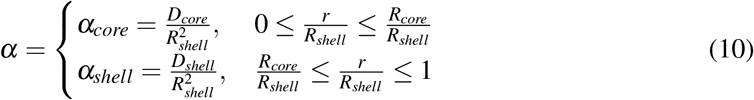

to scale the diffusion coefficient by the outer radius and yield a spatially dimensionless form. Note that the time dimension is still included. As defined in Equation (10), *α* is a piece-wise constant function.

In a technique called the method of lines [26, 27], the partial differential equation (PDE) in Equation (2) with *α* substituted is transformed into a system of ordinary differential equations (ODEs) by discretizing the normalized spatial domain and using a finite differencing scheme to approximate the spatial derivatives at the discrete points in the domain. The resulting ODEs can be solved using any built-in MATLAB ODE solver; here, we used ode45. The classic, second-order-accurate central finite difference discretization scheme in spherical coordinates with uniform grid spacing Δ*r* is [10, 28–31]

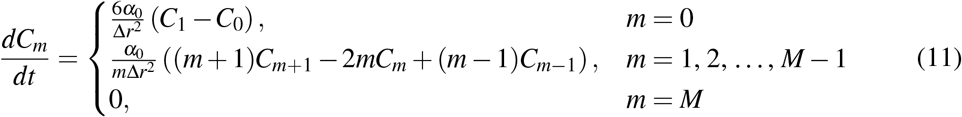

where *C*_*m*_(*t*) is the numerical approximation to *C*(*r*_*m*_,*t*) at the grid point *r*_*m*_ = *m*Δ*r* for *m* = 0, 1, …, *M* inside the normalized and discretized spherical domain with continuous time *t, M* is the number of discretizations in the domain, and Δ*r* = 1*/M*. Equation (11) holds for a constant value 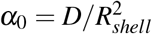 that is uniform throughout a spherical domain. Ford Versypt and Braatz [31] generalized Equation (11) to allow for any function *α*(*r,t*), including a piecewise constant function like Equation (10). However, that technique assumed that the grid spacing Δ*r* does not change. A fixed value of Δ*r* could be implemented for select values of *R*_*core*_ and *R*_*shell*_; however, here it is desirable to make the algorithm work for arbitrary combinations of *R*_*core*_ and *R*_*shell*_ to enable exploration of the design space of these sizes. Additionally, the interface boundary condition in Equations (4)–(5) needs to be satisfied. Thus, we adapted Equation (11) as the following schemes in the two layers for solving Equations (2)–(5):

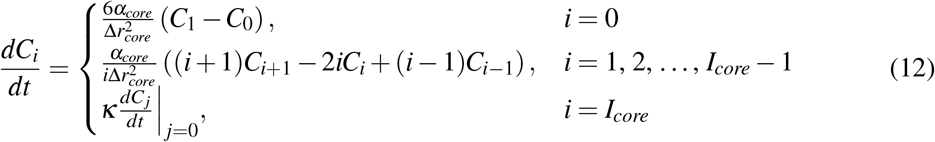

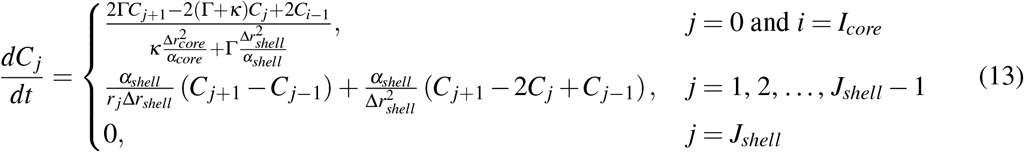

where *C*_*i*_(*t*) is the numerical approximation to *C*(*r*_*i*_,*t*) at the grid points *r*_*i*_ = *i*Δ*r*_*core*_ for *i* = 0, 1, …, *I*_*core*_ inside the core for 0 ≤ *r/R*_*shell*_ ≤ *R*_*core*_*/R*_*shell*_, Δ*r*_*core*_ = 1*/I*_*core*_ is the normalized grid spacing in the core, *I*_*core*_ is the number of spatial discretizations in the core, *C*_*j*_(*t*) is the numerical approximation to *C*(*r*_*j*_,*t*) at the grid points *r* _*j*_ = *I*_*core*_Δ*r*_*core*_ + *j*Δ*r*_*shell*_ for *j* = 0, 1, …, *J*_*shell*_ inside the shell for *R*_*core*_*/R*_*shell*_ ≤ *r/R*_*shell*_ *< R*_*shell*_*/R*_*shell*_, Δ*r*_*shell*_ = 1*/J*_*shell*_ is the normalized grid spacing in the shell, *J*_*shell*_ is the number of spatial discretizations in the shell, and

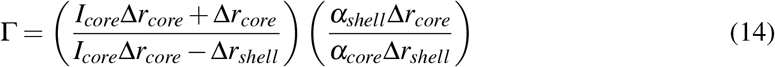

The schemes for the interior points in the shell layer (*j* = 1, 2, …, *J*_*shell*_ − 1) and at the interface (*i* = *I*_*core*_ and *j* = 0) and the selection of grid spacing and relationships between the grid spacing in the two layers to maintain second-order accuracy are derived in Section S1 in the Supplementary Material using techniques from [10, 32–34]. Figure S1 in the Supplementary Material shows a schematic representation of the core and shell interior points, the interface location, and the application of the fictitious node method at the interface.

Numerical techniques were used to evaluate the volume integrals in Equation (7) with discrete values of 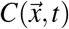 from the MATLAB finite difference solution. The MATLAB function `simps` [35] approximates each volume integral using Simpson’s method for numerical integration. In spherical coordinates, the volume integral becomes

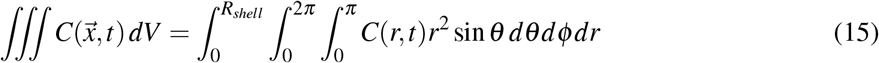

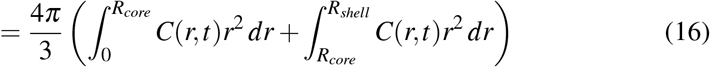

The final term allows for different grid spacing in the two layers and for non-uniform initial values *C*(*r*, 0) from Equation (6).

#### 2.3.2. Finite element approach in COMSOL Multiphysics

COMSOL Multiphysics was used to solve the model equations using the finite element method. The model geometry is a 2D axisymmetric structure, which reduces the 3D problem to a 2D problem by leveraging the symmetry about the vertical axis in the 2D plane. A 2D representation was selected instead of a 1D version as we wanted to generalize to other 2D axisymmetric (and 3D) geometries in the future. The solution procedure involves discretizing the spatial domain into simple geometric elements that are interconnected at common points with two or more elements. This process results in a system of algebraic equations that is solved simultaneously. A triangular mesh is used to define the 2D elements in the computational domain. In our implementation, the maximum element size allowed is 0.1 mm, while the minimum element size allowed is 2.4 × 10^*−*5^ mm based on the work of Van Kampen et al. [36], which also used COMSOL for modeling sustained intravitreal delivery from a DDS. The COMSOL interface “Transport of Diluted Species” is used as the physics package in the software to set up the PDE in Equation (2) to specify the boundary and initial conditions described in Section 2.2, and solve for the time-dependent drug concentration values in the domain. The drug transport equations are defined separately for each layer present in the DDS. At the interface between the layers, a “Partition Condition” boundary condition is used to specify κ.

Numerical techniques were used to evaluate the volume integrals in Equation (7) with discrete values of 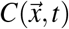 from the COMSOL finite element solution. The COMSOL built-in option `Non-local coupling: integration` was used to evaluate the volume integral with the spatially varying concentration profile of 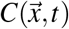 over the 2D axisymmetric domain that represents a microsphere.

The finite element approach in COMSOL Multiphysics and the finite difference approach in MATLAB were compared under the same parameter conditions. The drug concentration results at different locations inside the microspheres were used for verification purposes and were in good agreement (Section S2 and Figure S2 in the Supplementary Material).

### 2.4. Sensitivity analysis

We used one-at-a-time sensitivity approaches in MATLAB and COMSOL and the global variance-based Sobol’s method [37–39] implemented in COMSOL to assess the sensitivity of the model output to the input parameters. A sensitivity matrix *S* is defined with the general form

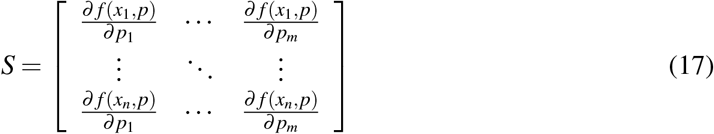

where *f* (*x*_*i*_, *p*) is a model output evaluated at input variable value *x*_*i*_ for *i* = 1, 2, …, *n* subject to the set of parameters *p* = *p*_1_, *p*_2_, …, *p*_*m*_. Generally, the input variables include the independent variable time *t*, the selected drug, and the geometric properties of the DDS: *R*_*core*_ and *R*_*shell*_. Here, we fixed *R*_*core*_ and *R*_*shell*_ to be the properties of the bi-layered microspheres from Jiang et al. [21] described in Section 2.1. For the model output of interest, we considered the cumulative release at the single time point of 28 days *Q*(28) from Equation (7) and the model solution of the concentration distribution from Equations (1)–(6) and the set of parameters *p* = {*B, D*_*core*_ = *D*_*Chi*_, *D*_*shell*_ = *D*_*PCL*_,*κ*}. The sensitivity analysis was performed for three regimes of diffusion coefficients in the two layers:

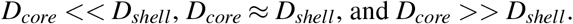

For the MATLAB local sensitivity analysis, finite differences were used to approximate the derivatives in Equation (17), and each parameter was increased one-at-a-time by 1%. Thus, the normalized local sensitivity *N*_*Q*(28,*p*)_ for the single output for each drug and four parameters is

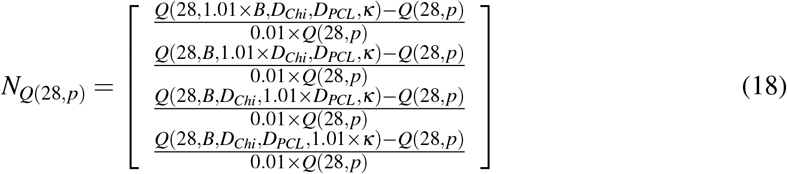

In COMSOL, the Morris one-at-a-time (MOAT) method [40–42] and Sobol’s method were both used from the uncertainty quantification module [43]; we allowed each parameter to be described by a normal distribution and a 1% standard deviation around the baseline value. The cumulative distribution function bounds were defined to constrain the parameter range within ±1% of the baseline value, with the lower bound set at 99% and the upper bound set at 101% of the baseline.

### 2.5. Parameter estimation

For the estimation of the parameters *p* = {burst release *B*, drug diffusion coefficients in the core *D*_*core*_ and the shell *D*_*shell*_, and the partition coefficient κ}, we used an ordinary least squares objective function with the general form:

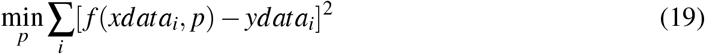

where *f* (*xdata*_*i*_, *p*) is the model output evaluated at values of input variables at which experimental measurements *ydata*_*i*_ were collected and *i* denotes discrete data points. Here, the data were from Jiang et al. [21], and the model outputs of interest were the cumulative release (Equation (7)) values at times *xdata*_*i*_ when the measurements for experimental cumulative drug release *ydata*_*i*_ were obtained. As in the sensitivity analysis, *R*_*core*_ and *R*_*shell*_ were fixed, and the parameter estimation process was conducted for both BSA and bevacizumab.

The MATLAB built-in function `lsqcurvefit` was used for estimating the parameters that minimized the least squares error (Equation (19)). The parameter values were normalized by dividing them by scaling factors specific to each parameter so that the maximum of the parameter range was set to 1. The estimated values were then multiplied by the scaling factors to yield the optimized parameter values. Tolerance was set to 0.001, and the maximum number of objective function evaluations for each parameter update was set to 800. All other optimization settings were at their default values.

We performed a preliminary parameter estimation described in Section S3 in the Supplementary Material over a broad parameter space that yielded feasible cumulative release profiles (Figure S3 and Table S1 in the Supplementary Material). The results from the preliminary parameter estimation (Figures S4 and S5 and Table S2 in the Supplementary Material) allowed us to reduce the parameter space of the problem before refining the parameter estimates. We performed a subsequent parameter estimation using a multi-start process with 50 initial guesses determined by Latin hypercube sampling of the parameter values bounded within the new limits in Table 1 with uniform sampling across the log of each parameter. The parameters with the lowest sensitivity were fixed. Upon completing the 50 multi-start parameter estimations, we compared the least squares error values for the parameter sets to which the `lsqcurvefit` algorithm converged, subject to each of the 50 initial parameter guesses. The parameter sets from the multi-start parameter estimations that yielded error values within 5% of the minimum error were deemed as acceptable. The resulting parameter values from the acceptable sets were averaged and then simulated for the average model results. Additionally, the parameters from the simulation with the lowest error value were considered the best model parameters. After comparing the best model and the average model, the average parameters were used in all further model simulations.

**Table 1:** Limits used in the multi-start parameter estimation.

| Parameter | Lower limit | Upper limit | Units |
| --- | --- | --- | --- |
| $B$ | 0 | 20 | % |
| $D_{Chi}$ | $1 \times 10^{-15}$ | $1 \times 10^{-13}$ | cm<sup>2</sup>/s |

$B$ : burst release. $D_{Chi} = D_{core}$ : drug diffusion coefficient in the chitosan core.

In COMSOL, we used the built-in `global least-squares objective` function in the optimization module to perform the parameter estimation (Equation (19)) for the four model parameters *p*. The parameter values were normalized by dividing them by scaling factors specific to each parameter; in COMSOL, these scaling factors were chosen such that the midpoint of the parameter range was set to 1. Tolerance was set to 0.001, and the maximum function evaluations was set to 1000.

### 2.6. Parameter uncertainty quantification

The sensitivity matrix *S* from Equation (17) was approximated using the finite difference approach in MATLAB with 1% perturbation in each of the parameters. The model output of interest *f* is the cumulative release *Q*(*t*) from Equation (7). The input variables *x*_*i*_ are discrete time points that correspond to 11 measurement times for cumulative release data from Jiang et al. [21]. The covariance of the cumulative release experimental data is collected in a matrix *ψ*. With *S* and *ψ*, the variance-covariance matrix *cov*(*p*) is approximated by [44]

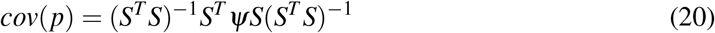

where *T* denotes the matrix transpose operation. The diagonal elements of this square matrix *cov*(*p*) give the variance of the parameters and were used to determine the parameters’ standard deviation. Finally, using this standard deviation in conjunction with the Student’s t inverse cumulative distribution function, the 95% confidence interval for each estimated parameter was calculated.

## 3. Results

### 3.1. Sensitivity analysis

We observed that the normalized sensitivity of the cumulative drug release at 28 days (Equation (18)) depended on the relative magnitudes of the diffusion coefficients in the chitosan core and the PCL shell (Figure 2). We identified three regimes for these relative magnitudes: *D*_*core*_ *<< D*_*shell*_, *D*_*core*_ ≈*D*_*shell*_, and *D*_*core*_ *>> D*_*shell*_ (Section S4 and Figure S6 in the Supplementary Material). For all the cases shown, the burst and the partition coefficient were tested at the same baseline values of 10 and 1, respectively. The MATLAB local sensitivity results are shown in Figure 2a,b,c, and the Sobol indices for the global sensitivity results from COMSOL are shown in Figure 2d,e,f. The sensitivity values from both approaches followed the same patterns regarding the ordering of the sensitivities of the four parameters; however, the insensitive parameters from the Sobol analysis had smaller magnitudes of the sensitivity indices than the corresponding values from the MATLAB local sensitivity analysis. Furthermore, the agreement between first-order and total Sobol indices across all three regimes suggested negligible interaction effects between parameters, indicating that each had an independent influence on cumulative release. These results were consistent with the results from the local MOAT analysis in COMSOL (Section S5 and Figure S7 in the Supplementary Material).

**Figure 2:**
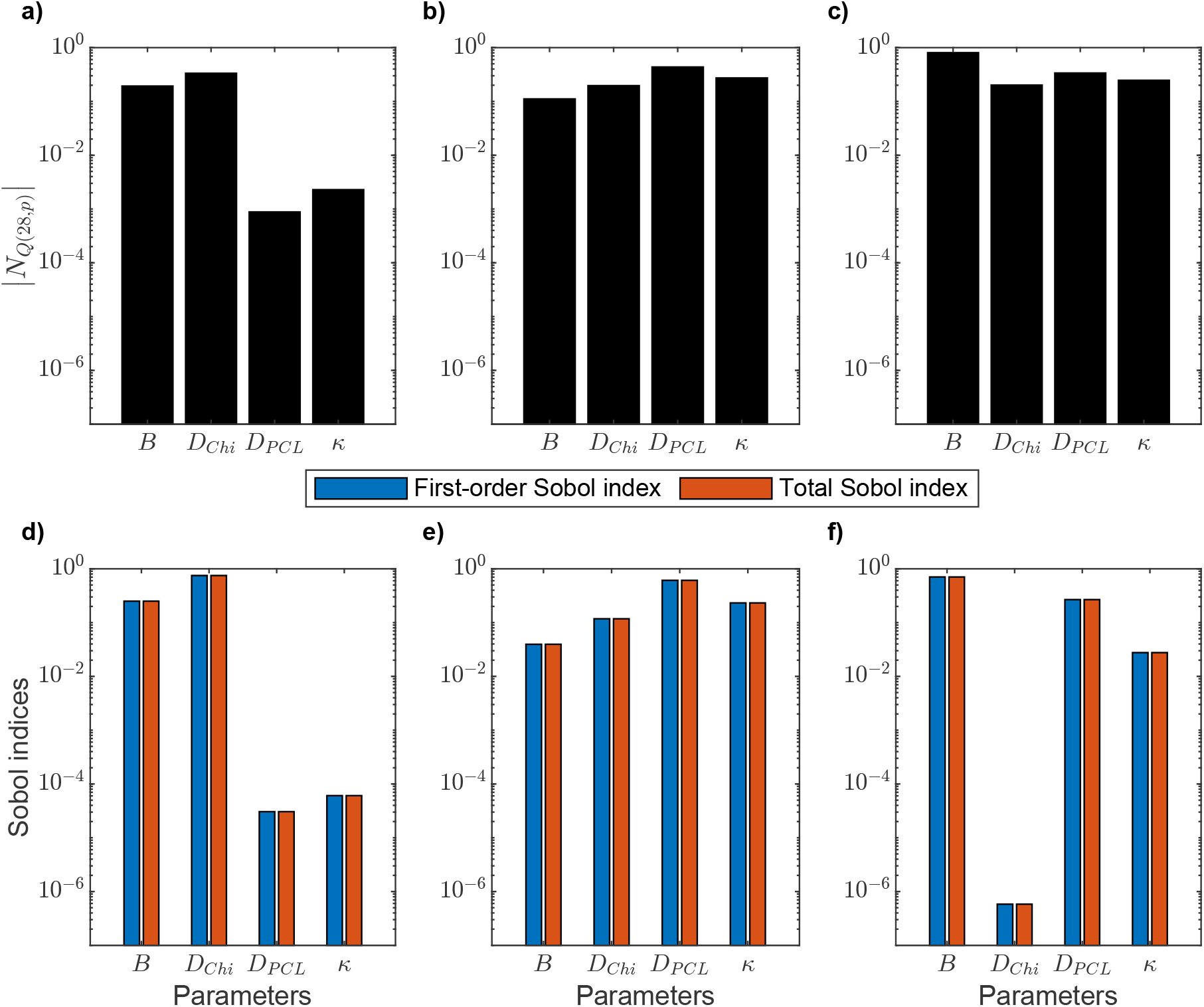
Sensitivity analysis by local and global methods for three regimes of the relative magnitudes of the diffusion coefficients in the chitosan core and the PCL shell. Top row: Absolute value of the normalized sensitivity of the cumulative drug release at 28 days |*N*_*Q*(28,*p*)_|obtained from the finite difference solution in MATLAB. Each parameter was varied one-at-a-time using a 1% increase. Bottom row: Sobol indices for the sensitivity analysis of parameters affecting cumulative drug release at 28 days obtained using the finite element solution in COMSOL. For each case, *B* = 10 and *κ* = 1. First column: *D*_*Chi*_ *<< D*_*PCL*_. Second column: *D*_*Chi*_ = *D*_*PCL*_. Third column: *D*_*Chi*_ *>> D*_*PCL*_. a) and d) *D*_*Chi*_ = 1 × 10^*−*15^ cm^2^/s and *D*_*PCL*_ = 1000 × *D*_*Chi*_ = 1 × 10^*−*12^ cm^2^/s, b) and e) *D*_*Chi*_ = 1 × 10^*−*14^ cm^2^/s and *D*_*PCL*_ = *D*_*Chi*_, and c) and f) *D*_*Chi*_ = 1 × 10^*−*12^ cm^2^/s and *D*_*PCL*_ = 0.001 × *D*_*Chi*_ = 1 × 10^*−*15^ cm^2^/s. *B*: burst release.κ: partition coefficient. *D*_*Chi*_ = *D*_*core*_: drug diffusion coefficient in the chitosan core. *D*_*PCL*_ = *D*_*shell*_: drug diffusion coefficient in the polycaprolactone shell.

The most influential parameter depended on the specific regime considered. When drug release was limited by drug diffusion in the chitosan core (*D*_*Chi*_ *<< D*_*PCL*_, Figure 2a,d), the drug diffusion coefficient in the chitosan core *D*_*Chi*_ was the most influential parameter with burst release *B* having a comparable effect. In contrast, the drug diffusion in the PCL shell *D*_*PCL*_ and the partition coefficient *κ* had negligible impacts on the cumulative drug release at 28 days. When the diffusion coefficients were the same (*D*_*Chi*_ = *D*_*PCL*_, Figure 2b,e), the sensitivity shifted, and the drug diffusion coefficient in the PCL shell *D*_*PCL*_ became the dominant parameter, followed by the partition coefficient *κ* and the drug diffusion coefficient in chitosan *D*_*Chi*_. In this case, the burst release *B* had the lowest impact on the cumulative drug release at 28 days. When drug release was limited by drug diffusion in the PCL shell (*D*_*Chi*_ *>> D*_*PCL*_, Figure 2c,f), the burst release *B* was the most sensitive parameter, followed by the drug diffusion coefficient in the PCL shell *D*_*PCL*_ and partition coefficient *κ*. In this regime, the drug diffusion coefficient in the chitosan core *D*_*Chi*_ had the smallest effect on the cumulative drug release at 28 days.

### 3.2. Parameter estimation

The preliminary parameter estimation considering all four parameters for both BSA and bevacizumab (Section S3 in the Supplementary Material) showed that *D*_*Chi*_ *<< D*_*PCL*_ best fit the chitosan-PCL core-shell microsphere DDS data used here (Figures S3 and S4 and Table S2 in the Supplementary Material). This parameter regime was insensitive to the parameters *D*_*PCL*_ and *κ* (Figure 2a,d), meaning that their values did not substantially influence the model. Therefore, we set *κ* = 1 and *D*_*PCL*_ = 1000 × *D*_*Chi*_ and only further refined the parameter estimates for *D*_*Chi*_ and *B*. This reduced model complexity and accelerated the parameter estimation and solution procedure without compromising accuracy.

All the error values after each completed parameter estimation for BSA release in MATLAB and COMSOL (Figure 3a,b, respectively) and for bevacizumab release (Figure 4a,b) were within a 5% error threshold and, therefore, were used when averaging the parameters. Thus, the parameter estimation process was insensitive to variations in initial guesses for this regime of diffusion coefficients. For both MATLAB and COMSOL, the average and best (minimum error over all the multi-start parameter estimations) model predictions overlapped and were close to experimental data (Figure 3c,d and Figure 4c,d), indicating that the implementation of the model in both software programs predicted the same BSA and bevacizumab release from the bi-layered microspheres in agreement with data. Furthermore, Table 2 shows the similarity between the average and best model parameters for each software program and the similarity between the best models from both software programs.

**Table 2:**
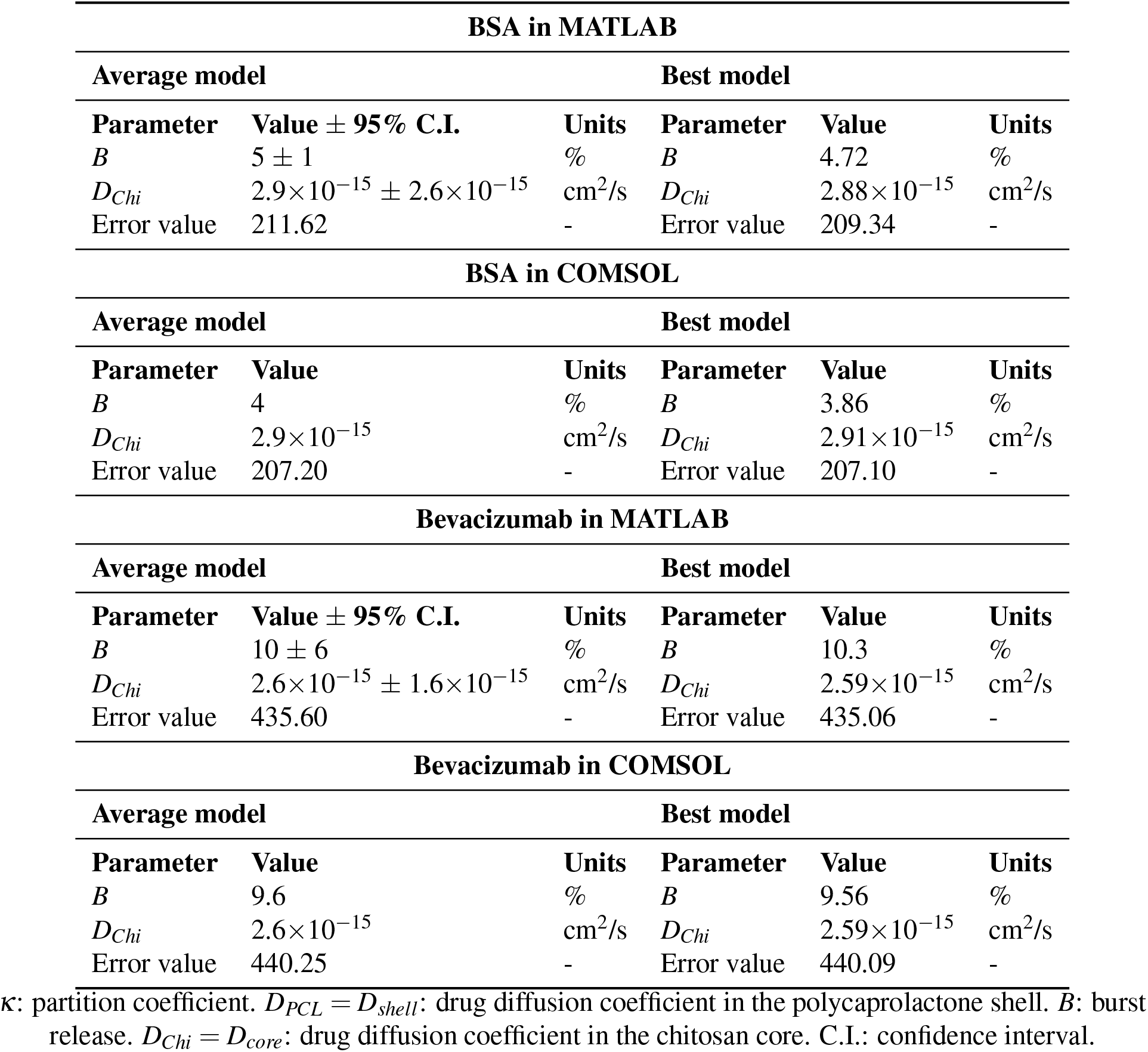
Estimated parameters for fitting the model to data for drug-loaded chitosan-PCL core-shell microspheres from Jiang et al. [21]. The average model was obtained by averaging the parameters obtained from all the optimization runs that achieved an error within 5% of the minimum error after 50 multi-start attempts. The best model corresponds to the parameter set with the lowest error value. The other model parameters were fixed as κ= 1 and *D*_*PCL*_ = 1000 *×D*_*Chi*_.

**Figure 3:**
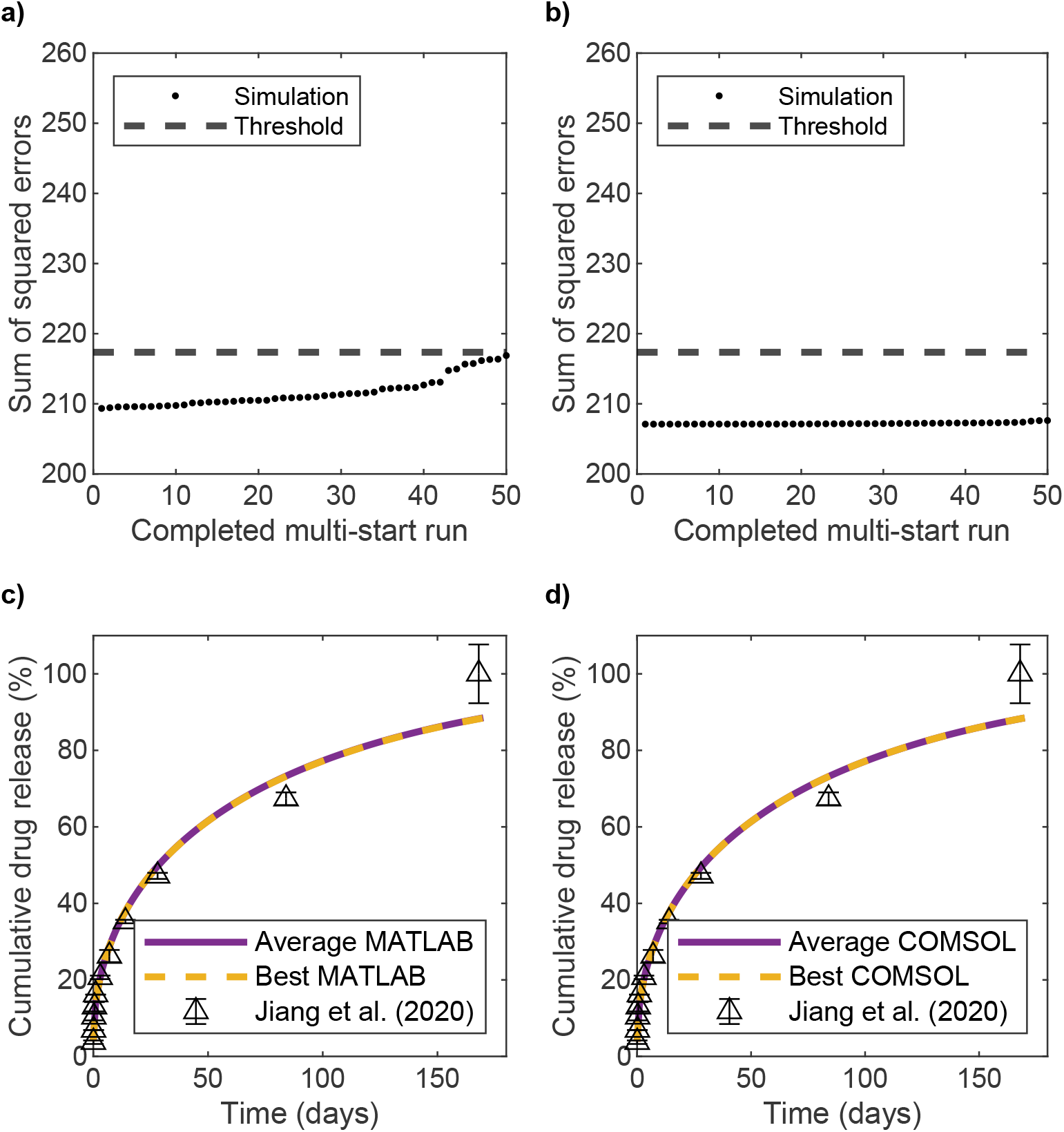
Error values and cumulative drug release profiles for BSA release after 50 multi-start parameter estimations. a) Error values in MATLAB. b) Error values in COMSOL. c) Average and best MATLAB models compared to experimental data. d) Average and best COMSOL models compared to experimental data. Experimental data from Jiang et al. [21].

**Figure 4:**
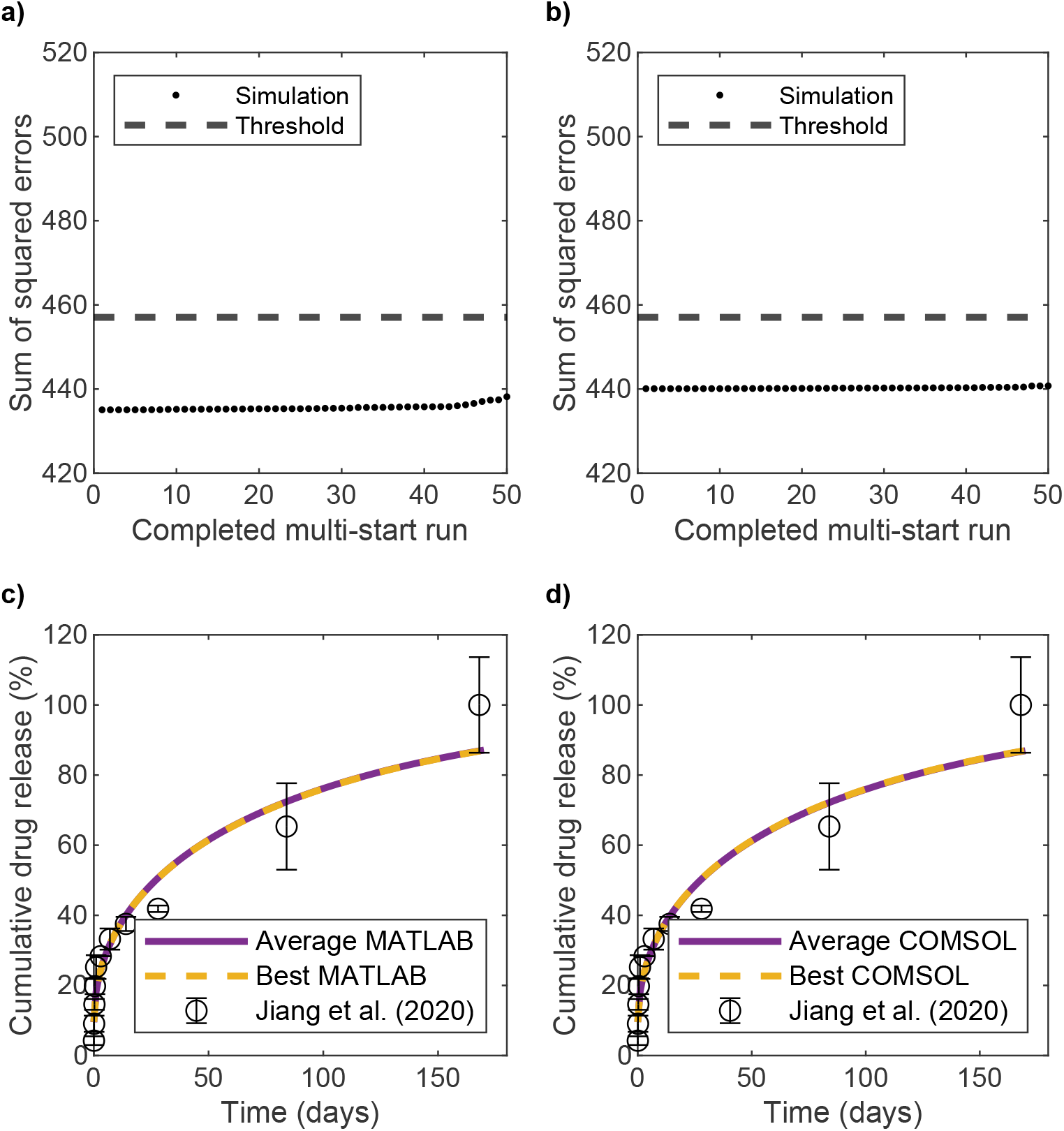
Error values and cumulative drug release profiles for bevacizumab release after 50 multi-start parameter estimations. a) Error values in MATLAB. b) Error values in COMSOL. c) Average and best MATLAB models compared to experimental data. d) Average and best COMSOL models compared to experimental data. Experimental data from Jiang et al. [21].

Table 2 also shows the 95% confidence intervals for the uncertainty in parameter values as calculated with the MATLAB results according to Section 2.6. For both BSA and bevacizumab, burst release had a relatively narrower confidence interval than that for the drug diffusion coefficient in the chitosan core. For both drugs, the confidence intervals did not cross into the negative regions for any parameter. Among the parameters estimated in COMSOL, none of them fell outside the 95% confidence intervals obtained from MATLAB.

### 3.3. Predictive capabilities

The computational model was used to explore the impact of the sizes of both chitosan and PCL layers on cumulative drug release and the drug release rate to optimize polymer layer sizes for bevacizumab release. The predicted bevacizumab release rate was obtained from MATLAB using Equations (8) and (9) for the case of *t > t*_0_. Alternatively, the release rate could be computed in COMSOL as the line integral of the total normal flux multiplied by the molecular weight of bevacizumab; when simulating microspheres of different sizes in COMSOL, the computed flux must also be scaled by a geometric factor that accounts for changes in surface area due to variations in sphere volume relative to the baseline size. The bevacizumab loaded amount *A*_*load*,0_ for all simulations was 1.03 mg. This loading amount is feasible and is representative of the range of 0.99–1.61 mg of BSA or bevacizumab loaded into 4 mg of chitosan or chitosan-PCL microspheres in Jiang et al. [21].

The target for the optimal design was to maintain the drug release rate above the estimated threshold as long as possible while ensuring that a substantial portion of the total drug was delivered based on cumulative release. The threshold for drug release was derived from a previously published study that established that an injection of 12.5 µg of bevacizumab was sufficient to reduce neovascularization leakage during one week in patients with proliferative diabetic retinopathy [45]. By dividing the injected amount over 7 days, the approximate threshold can be rounded to 2.0 µg/day. This is consistent with Carichino et al. [46] in which the authors calculated that a retinal anti-VEGF production rate of 1.12–2.62 µg/day would be as effective as an injection of ranibizumab (another therapeutic that slows neovascularization). Here, the cumulative drug release threshold was set at 90%, as it is representative of practical drug release scenarios, and this threshold was also used in another study of sustained intravitreal protein delivery from a DDS [36].

The results in this section were simulated in MATLAB using the average model parameters for bevacizumab in Table 2. Increasing the radius of the chitosan core (left to right in Figure 5) resulted in slower cumulative drug release due to the greater distance the drug had to diffuse through the polymer. A similar but weaker effect was observed when increasing the PCL shell layer (top to bottom in Figure 5). A large increase in the PCL shell would be required to slow the cumulative drug release substantially, consistent with the model’s insensitivity to *D*_*PCL*_ in the parameter regime where the chitosan core controls the release (*D*_*core*_ *<< D*_*shell*_). For the fastest cumulative drug release (top left in Figure 5), the 90% cumulative release threshold was achieved between 12–13 days, whereas for the slowest cumulative drug release (bottom right) only about 70% of the loaded drug was released by 6 months. We extended the simulation time to 360 days (Figure S8 in the Supplementary Material), and the slowest case increased to 85% cumulative release by that time.

**Figure 5:**
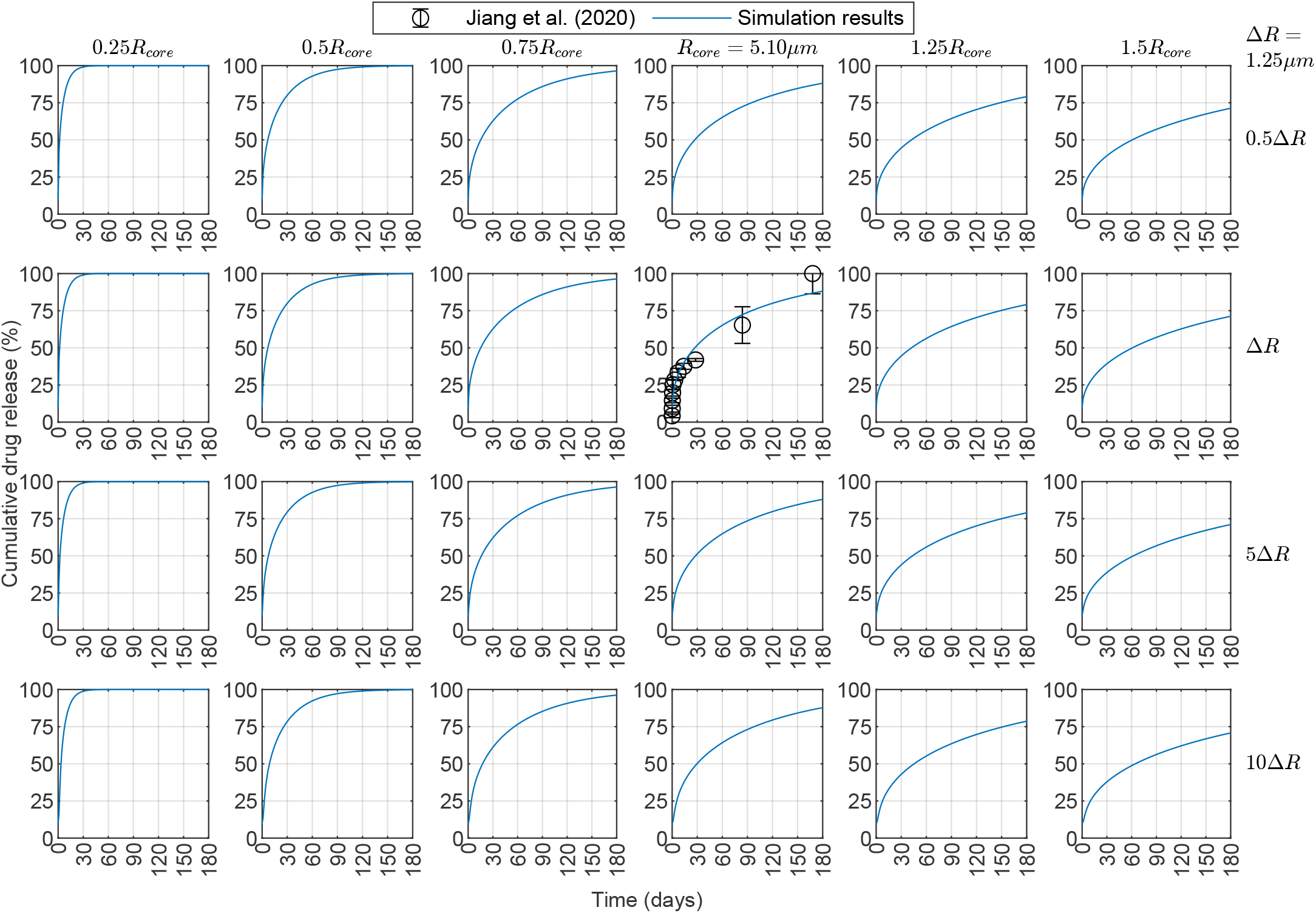
Cumulative drug release profiles for bevacizumab released from core-shell microspheres with drug loaded in the core only. Results were obtained in MATLAB with *D*_*Chi*_ = 2.6 × 10^*−*15^ cm^2^/s, *D*_*PCL*_ = 2.6 × 10^*−*12^ cm^2^/s, *B* = 10%, and *κ* = 1 (average model parameters from Table 2). Experimental data for *R*_*core*_ = 5.10 µm and Δ*R* = *R*_*shell*_ − *R*_*core*_ = 1.25 µm are from Jiang et al. [21]. Each panel shows profiles for different chitosan-PCL configurations where the core radius and shell thickness are varied by multipliers to the baseline dimensions *R*_*core*_ and Δ*R* considered from Jiang et al. [21]. The core radii are labeled on the columns, and the shell thicknesses are labeled on the rows. *D*_*Chi*_ = *D*_*core*_: drug diffusion coefficient in the chitosan core. *D*_*PCL*_ = *D*_*shell*_: drug diffusion coefficient in the polycaprolactone shell. *B*: burst release. *κ*: partition coefficient.

Because of the assumption that *C*_*core*,0_ is the same for all microspheres, those with larger *R*_*core*_ were loaded with more drug; fewer of these microspheres were considered in a dose to maintain the same *A*_*load*,0_ for all simulations. Reducing the chitosan core radius below the baseline resulted in a faster release, shortening the duration of the therapeutic delivery rate (Figure 6). In contrast, increasing the chitosan radius beyond 1.5 times its baseline value was not substantially beneficial for the drug release rate. Furthermore, variations in the PCL layer had little effect on the drug release rate within the parameter regime that best fits the data, and a substantial increase in the PCL layer would be required to observe any appreciable change in the release rate. For instance, for baseline chitosan and PCL, a 10-fold increase of the PCL layer thickness only extended the time above the target 2 µg/day drug release rate from 111 days to 114 days. This limited PCL layer dependence was not surprising given that no drug was loaded into the PCL layer and *D*_*core*_ *<<D*_*shell*_. Additionally, considering a ±10% window around the release rate threshold gave a range of days at which the window of the minimum therapeutic release threshold was reached; the duration of this therapeutic target range increased with the chitosan radius (green shaded region in Figure 6). Finally, all microspheres with chitosan radius at the baseline value or larger maintained their drug release rate over 1 µg/day by the end of the simulations at 180 days (Figure 6). We extended the simulation time to 360 days and lowered the drug release rate threshold to 1 µg/day (Figure S9 in the Supplementary Material), and all the chitosan-PCL configurations considered reached that threshold by 240 days.

**Figure 6:**
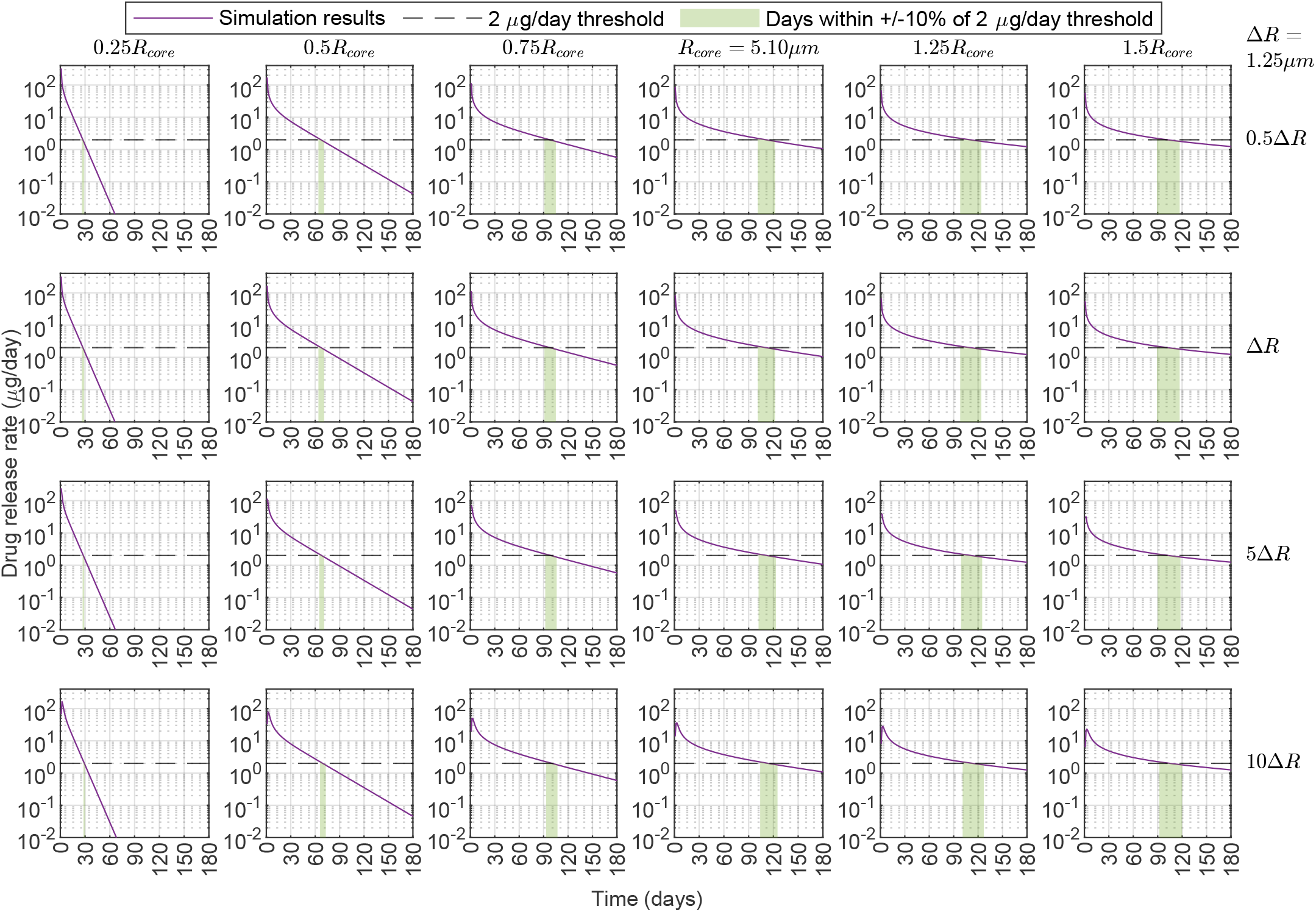
Drug release rate profiles for bevacizumab released from core-shell microspheres with drug loaded in the core only. Results were obtained in MATLAB with D*_Chi_* = 2.6×10^15^ cm^2^/s, D*_PCL_* = 2.6×10^12^ cm^2/^/s, *B* = 10%, and *κ* = 1 (average model parameters from Table 2). Each panel shows profiles for different chitosan-PCL configurations where the core radius and shell thickness are varied by multipliers to the baseline dimensions *R_core_* = 5.10 μm and Δ *R* = *R_shell_* − *R_core_* = 1.25 μm considered from Jiang et al. [21]. The core radii are labeled on the columns, and the shell thicknesses are labeled on the rows. On each panel, the purple curve is the simulation results, the dashed line shows the 2 μg/day drug release rate threshold, and the shaded green region highlights the days within ± 10% of the 2 μg/day drug release rate threshold. D*_Chi_* = D*_core_*: drug diffusion coefficient in the chitosan core. D*_PCL_* = D*_shell_*: drug diffusion coefficient in the polycaprolactone shell. *B*: burst release. *κ*: partition coefficient.

The time to reach the 90% cumulative release threshold increased monotonically with *R*_*core*_ and Δ*R*, with a stronger effect of *R*_*core*_ (Figure 7a). The duration of time above the 2 µg/day release rate threshold reached a maximum around 1.2 times the baseline chitosan radius (Figure 7b); variations in the PCL layer thickness had little effect. Figure 7c shows the surface plots for threshold cumulative drug release and drug release rate from Figure 7a and Figure 7b to visualize their intersection. Figure 7d shows two of the three axes of Figure 7c with the largest and smallest Δ*R* baseline mul-tipliers plotted. The shading around the curves indicates the intervals where the release rate was within ±10 % of the target 2 µg/day. While the surface plot in Figure 7b went through a maximum, Figure 7d shows that values of ≈1–1.4 × *R*_*core*_ had similar numbers of days above the drug release rate threshold, particularly when considering the intervals around the target threshold.

**Figure 7:**
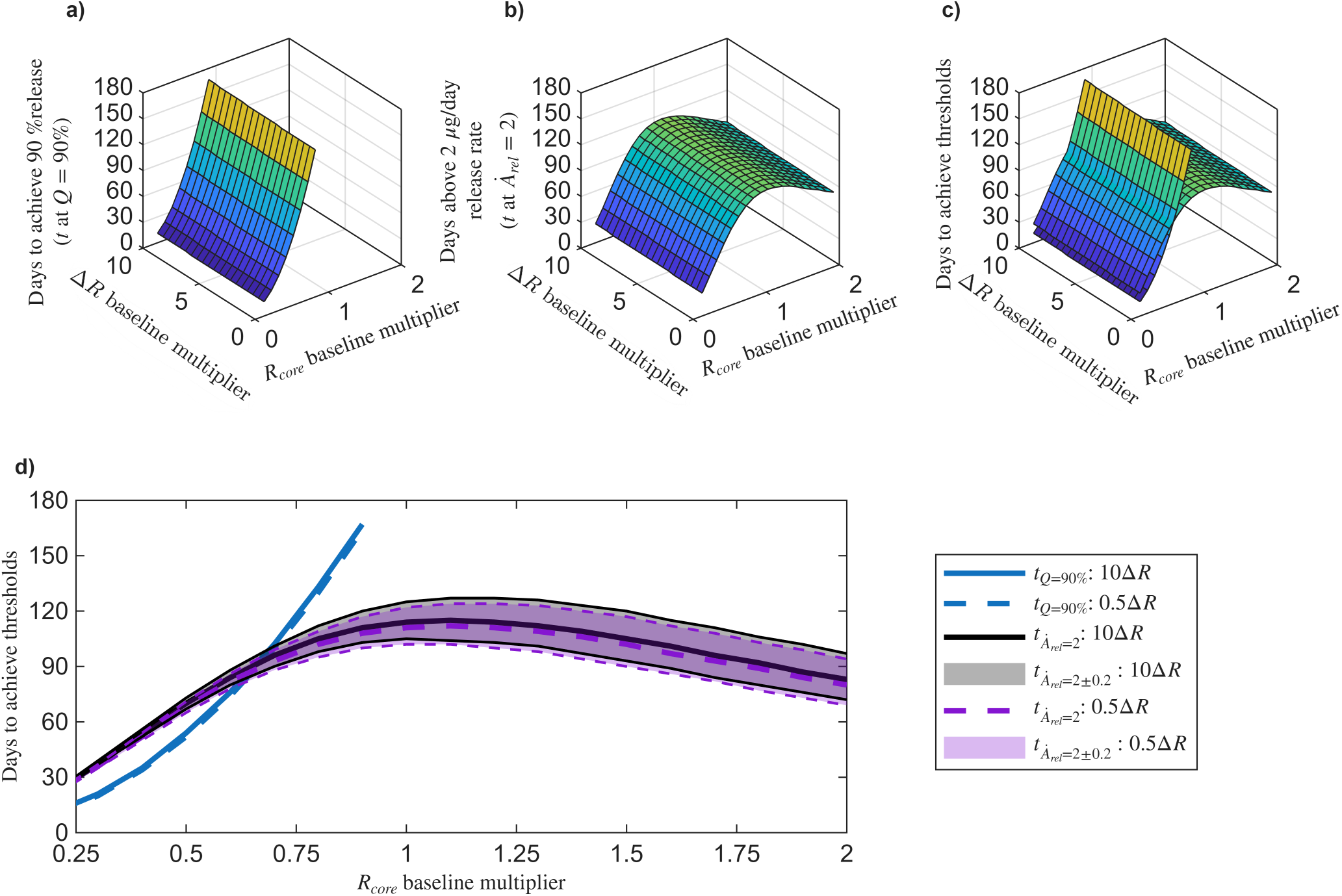
Time in days to reach cumulative release (*Q*) and release rate (*Ȧ*_*rel*_) thresholds for different chitosan-PCL configurations (including those from Figures 5 and 6) where the core radius and shell thickness are varied by multipliers to the baseline dimensions *R*_*core*_ = 5.10 µm and Δ*R* = *R*_*shell*_ −*R*_*core*_ = 1.25 µm considered from Jiang et al. [21]. a) Time in days to reach cumulative release threshold of *Q* = 90% as a function of *R*_*core*_ and Δ*R* baseline multipliers. b) Time in days to reach release rate threshold of *Ȧ*_*rel*_ = 2 µg/day as a function of *R*_*core*_ and Δ*R* baseline multipliers. c) 3D view of the surfaces from panels a) and b) combined. d) 2D view of panel c) in the projection onto the time vs. *R*_*core*_ baseline multiplier axes. Δ*R* baseline multipliers of 0.5 and 10 are shown in panel d) along with shaded regions denoting the intervals where the release rates are within *±* 10% of the threshold. Results for all panels were obtained in MATLAB with *D*_*Chi*_ = 2.6 *×* 10^*−*15^ cm^2^/s, *D*_*PCL*_ = 2.6 *×* 10^*−*12^ cm^2^/s, *B* = 10%, and *κ* = 1 (average model parameters from Table 2). *R*_*core*_ = 5.10 µm and Δ*R* = *R*_*shell*_ *− R*_*core*_ = 1.25 µm are from Jiang et al. [21]. *D*_*Chi*_ = *D*_*core*_: drug diffusion coefficient in the chitosan core. *D*_*PCL*_ = *D*_*shell*_: drug diffusion coefficient in the polycaprolactone shell. *B*: burst release. *κ*: partition coefficient.

One possible optimal design was the chitosan-PCL configuration that achieved the intersection of the time to reach the cumulative drug release and drug release rate thresholds (Figure 7c,d). This occured for a chitosan-PCL bi-layered microsphere configuration with ≈0.65 × *R*_*core*_ and 1xΔ*R* (as there was not substantial dependence on the PCL layer thickness). This configuration sustained a drug release rate over 2 µg/day and achieved 90% drug depletion within 90 days, minimizing the time between falling below the therapeutic release rate and fully depleting the microspheres of their drug payloads. An alternative definition of an “optimal” chitosan-PCL configuration was where the time above the drug release rate threshold reached a maximum, as long as the microspheres had not surpassed the 90% cumulative release threshold. For the release rate threshold of 2 µg/day for more than 100 days, this optimal chitosan-PCL bi-layered microsphere configuration was ≈1–1.4 × *R*_*core*_ (ideally, the largest size in this range is the preferred option because it held more drug payload) and 1xΔ*R*. The tradeoff with this design is that the microspheres had released less than 90% of their load by the time that they fell below the therapeutic release rate threshold.

If we instead considered the therapeutic drug release rate threshold to be 1 µg/day, then the intersection of the time to reach the cumulative drug release and drug release rate thresholds (Figure S10c,d in the Supplementary Material) occurred around 180 days for a chitosan-PCL bi-layered microsphere configuration with ≈0.95 × *R*_*core*_ and 1xΔ*R*. The time above the drug release rate threshold reached a broad maximum region (Figure S10d in the Supplementary Material) around 220 days at ≈1.4–1.9 × *R*_*core*_ and 1xΔ*R*. While these configurations had not reached 90% cumulative release, they had exceeded 75% release by this time (Figure S8 in the Supplementary Material).

To facilitate the design and analysis of core–shell drug delivery systems, we developed an interactive MATLAB live script that enables rapid exploration of how the dimensions of the polymer layers and transport parameters influence the dynamics of drug release (Figure 8). The tool accepts user-defined inputs for baseline core radius and shell thickness, along with the minimum, maximum, and step sizes for the multipliers used to scale these baseline values. Additional input parameters include maximum simulation time, thresholds for both cumulative drug release and daily release rate, and a tolerance range around the release rate threshold to allow for flexible performance assessments. Users can also modify the values of previously estimated parameters, including drug diffusion coefficients, partition coefficient, and burst release, to examine the impacts of these parameters on release profiles. The tool generates plots corresponding to several possible combinations of core and shell size multipliers within the specified range. Each plot displays either cumulative release (similar to Figure 5) or release rate (similar to Figure 6) as a function of time, enabling visualizations of how drug delivery performance changes with geometry specifications. This flexible and user-friendly framework allows researchers to efficiently assess design trade-offs and optimize core–shell configurations for long-term therapeutic delivery.

**Figure 8:**
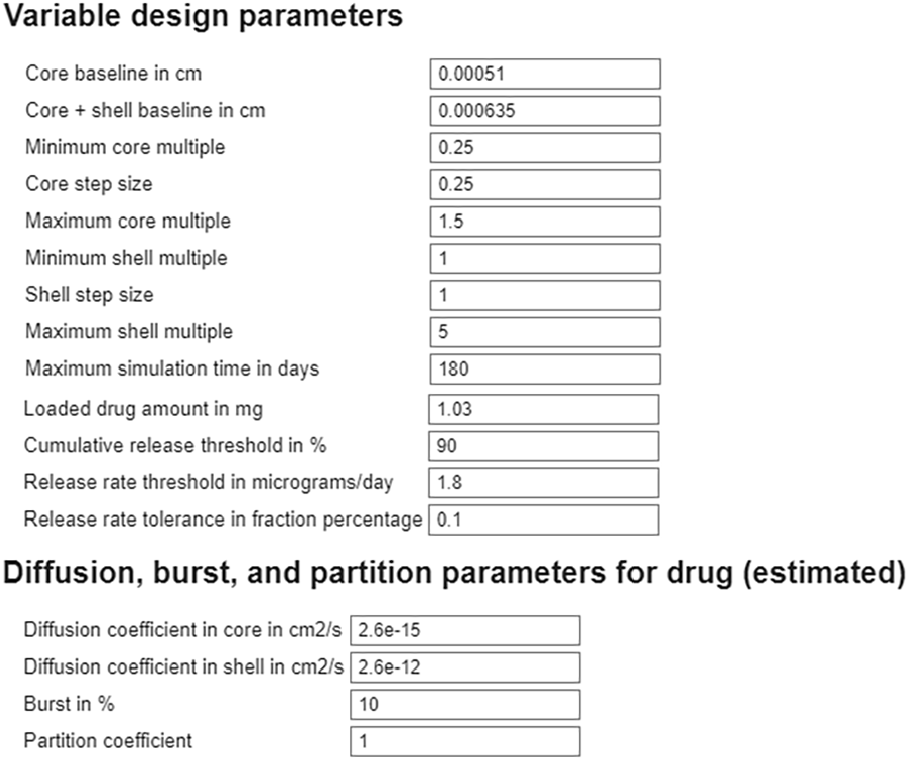
Script screenshot of the MATLAB-based simulation tool for core-shell drug delivery systems (DDSs). Users can specify baseline size values, range of multipliers for those baseline values, thresholds, and tolerances. The tool also allows modification of estimated transport parameters. The output consists of a drug cumulative release plot and a drug release rate plot for all combinations of core and shell dimensions, facilitating the design and optimization of long-acting DDSs.

## 4. Discussion

In this work, we combined mathematical modeling, numerical simulations, and sensitivity-guided parameter estimation into a computational framework tailored for chitosan-PCL bi-layered microspheres for intravitreal controlled release to treat chronic retinal diseases. We developed and parameterized a diffusion-controlled model to characterize long-term drug release from core-shell microspheres, focusing on intravitreal therapies for chronic diseases like wet AMD, where repeated injections present challenges to patient compliance and safety. Using experimental release data for BSA and bevacizumab, we estimated key transport parameters (drug diffusion coefficients, partition coefficient, and the burst release) using both finite difference (MATLAB) and finite element (COMSOL) implementations.

While prior models [19, 20] have applied similar frameworks to glucose and metronidazole release from multi-layered systems, they often made simplifying assumptions, such as unity partition coefficients, no burst release, and short delivery time frames. In contrast, our model explicitly incorporates burst release and partitioning effects (albeit with this effect optimized to a value of 1), extending the analysis to clinically relevant long-term delivery periods and offering a more comprehensive representation of ocular drug transport from bi-layered spherical DDSs.

We verified our computational framework through strong agreement between MATLAB and COMSOL implementations (Figures 3 and 4 and Figure S2 in the Supplementary Material). The 50 multi-start parameter estimations in MATLAB (using parallel `parfor loops`) required 12–16 days to complete on a Windows PC with 32 GB RAM and an 11th Gen Intel core i7-11700 processor with eight cores. In comparison, COMSOL estimations took around 6 days and required manual adjustments to initial guesses. Despite slight differences in the estimated values for the burst release and the diffusion coefficients of drugs in chitosan (Table 2), both implementations yielded consistent results overall. The MATLAB model was used for design exploration due to its faster runtime for the forward simulation problem (as compared to the inverse parameter estimation problem) and better support for automation. In contrast, COMSOL offers advantages that could be leveraged to deal with more complex geometries where simple approximations are insufficient.

The sensitivity analysis identified three distinct diffusion regimes and quantified how each parameter influenced the 28-day cumulative drug release (Figure 2). The relative importance of the parameters in the sensitivity analysis depended on the specific regime considered. In the core-limited regime (*D*_*core*_ *<< D*_*shell*_), changes in core diffusivity had the greatest impact, while in the shell-limited regime (*D*_*core*_ *>> D*_*shell*_), diffusivity in the shell was more influential than diffusivity in the core; this is consistent with the findings from Barchiesi et al. [20]. Notably, both local and global sensitivity analyses ranked parameters consistently across the three regimes and showed minimal interaction effects.

To demonstrate the utility of our framework, we applied it to a case study involving chitosan–PCL microspheres loaded with bevacizumab, a widely used anti-VEGF therapeutic. Using clinically relevant thresholds of 2 µg/day [46] for daily delivery and 90% cumulative release [36], we examined how varying the core radius (*R*_*core*_) and shell thickness (Δ*R*) influenced therapeutic performance. While increasing *R*_*core*_ or Δ*R* slowed cumulative drug diffusion (Figures 5 and 7a), an increase in *R*_*core*_ from its baseline value had little effect on drug release rate until it exceeds 1.5 × *R*_*core*_ where the release rate began to decline (Figures 6 and 7b). Conversely, reducing *R*_*core*_ accelerated drug release, shortening the duration above the release rate threshold. Δ*R* increase needed to be substantial to observe a slight increase on the days the drug release rate stayed above the threshold. This implies that tuning *R*_*core*_ is the best way to optimize the therapeutic delivery and that Δ*R* adjustments need to be substantial for meaningful drug release modifications.

For a therapeutic release rate target of 2 µg/day of bevacizumab, the best configuration for sustained therapeutic release and efficient drug depletion was around 0.65 × *R*_*core*_ and 1 × Δ*R* at 90 days (Figure 7c,d). The alternative best configuration that maximized the time above the therapeutic release rate threshold was around 1.4 × *R*_*core*_ and 1 × Δ*R* (Figure 7c,d) for 100 days, while leaving some loaded drug to release at a sub-therapeutic daily rate for a longer time. For a reduced therapeutic release rate target of 1 µg/day of bevacizumab, the equivalent best configurations were for 0.95 × *R*_*core*_ and 1 × Δ*R* at 180 days and 1.9 × *R*_*core*_ and 1 × Δ*R* for 220 days (Figure S10c,d in the Supplementary Material). These findings highlight the importance of prioritizing tuning the core size in DDS design in the core-limited regime (*D*_*core*_ *<< D*_*shell*_).

All computational models have their limitations. Here, we have only parameterized the models to *in vitro* data. The design trends should still hold when moving *in vivo*, but the actual days to reach the release thresholds will be influenced by the *in vivo* conditions. Notably, erosion likely should be considered in that case as PCL enzymatic degradation *in vivo* was shown to be faster than hydrolytic degradation *in vitro* [23, 47]. We have also assumed perfect sink external boundary conditions, where the released drug is instantaneously removed from the DDS surface. The model defined here could be combined with a more realistic treatment of the intravitreal conditions, such as in our recent three-dimensional ocular pharmacokinetics models for rabbit and human eyes [48]. Together, the intravitreal drug release could be validated or re-parameterized using *in vivo* data as it becomes available.

Finally, an additional contribution of this work is the development of a generalizable and open computational framework for designing and optimizing diffusion-controlled core-shell DDSs. Our dual implementation in MATLAB and COMSOL enables experimentalists and modelers to estimate drug-specific parameters and validate models against release data, demonstrated here for the relatively simple bi-layered microsphere geometry. The COMSOL code can be easily modified to accommodate more complex geometries. Moreover, our MATLAB design script facilitates the efficient exploration of design trade-offs. The framework of this work supports model-informed DDS design, accelerating the development of long-acting ocular therapeutics. We have shared all of these codes openly at https://github.com/ashleefv/LayeredSpheres_DDSdesign [49].

## 5. Conclusions

In this study, we developed a diffusion-based computational model to investigate long-acting intravitreal drug delivery using core–shell microspheres. The model accounts for key transport process parameters, including burst release, drug partitioning, and diffusion coefficients across bilayered polymeric spheres. Sensitivity analyses revealed different parameter dominance depending on the specific parameter regimes driving drug release. In the chitosan-PCL case study, we identified drug diffusivity in the core and burst release as the primary drivers of the long-term release behavior. The estimated transport parameters were fitted to experimental release data for BSA and bevacizumab using both finite difference (MATLAB) and finite element (COMSOL) methods. Using clinically relevant thresholds for therapeutic delivery, we simulated drug release from chitosan–PCL microspheres and showed that core radius has a greater impact on therapeutic performance than shell thickness in the data-driven parameter regime. We further developed a customizable MATLAB tool to explore the trade-off of using different layer sizes through automated simulations, providing visual outputs of the profiles of cumulative drug release and drug release rate across a range of core and shell sizes. This tool supports rapid hypothesis testing and formulation design, and the generalizable modeling framework enables easy adaptation to other bi-layered spherical DDSs. Our findings underscore the importance of mechanistic modeling in informing the design of sustained-release drug delivery systems to enhance the treatment of chronic ocular diseases.

## Data availability

No new data were created or analyzed in this study. We have provided our codes for MATLAB 2024b and COMSOL v6.2 in a repository at https://github.com/ashleefv/LayeredSpheres_DDSdesign [49].

## Acknowledgments

This work was supported by National Institutes of Health grants R35GM133763 to ANFV and R01EB032870 to KESR and ANFV, an Owen Locke Foundation grant to KESR, and the University at Buffalo. We thank our research group members for their thorough feedback on this manuscript and for their helpful discussions.

