## Supplementary Material for "Modeling and Design of Chitosan-PCL Bi-Layered Microspheres for Intravitreal Controlled Release"

### S1. Derivation of finite difference schemes

#### *S1.1. Interior nodes*

The finite difference equations for diffusion in a sphere with the inner and outer boundary conditions defined in Section 2 are well-known and are shown in Equations (12) and (13) for $i = 0$ and $j = J_{shell}$, respectively. The finite difference scheme for the interior nodes of the core layer is also in a standard form for $i = 1, 2, \ldots, I_{core} - 1$ in Equation (12). Here, we first rewrite the interior form to consider how the radial position impacts the scheme, generalizing it to the case where the grid spacing is not uniform and the indexing restarts in the shell layer. Let $\hat{r}$ be the radial position from the center of the sphere and $r$ be the normalized radial position, where $r = \hat{r}/R_{shell}$. The diffusion equation in dimensionless spherical coordinates using $\alpha$ from Equation (10) and expanding the derivative in Equation (2) gives

$$\frac{\partial C}{\partial t} = \begin{cases} \dfrac{2\alpha_{core}}{r}\dfrac{\partial C}{\partial r} + \alpha_{core}\dfrac{\partial^2 C}{\partial r^2}, & 0 \le r \le R_{core}/R_{shell} \\[2ex] \dfrac{2\alpha_{shell}}{r}\dfrac{\partial C}{\partial r} + \alpha_{shell}\dfrac{\partial^2 C}{\partial r^2}, & R_{core}/R_{shell} \le r < 1 \end{cases} \tag{S1}$$

[1]Current address: School of Aerospace and Mechanical Engineering, The University of Oklahoma, Norman, OK, USA

Applying centered finite differences for $0 < r < R_{core}/R_{shell}$ gives

$$\frac{dC_i}{dt} = \frac{\alpha_{core}}{r_i \Delta r_{core}}\left(C_{i+1} - C_{i-1}\right) + \frac{\alpha_{core}}{\Delta r_{core}^2}\left(C_{i+1} - 2C_i + C_{i-1}\right), \quad i = 1, 2, \ldots, I_{core} - 1 \tag{S2}$$

which can be rewritten as

$$\frac{dC_i}{dt} = \frac{\alpha_{core}}{\Delta r_{core}^2}\left(\frac{r_{i+1}}{r_i}C_{i+1} - 2\frac{r_i}{r_i}C_i + \frac{r_{i-1}}{r_i}C_{i-1}\right), \quad i = 1, 2, \ldots, I_{core} - 1 \tag{S3}$$

In the core, $r_i = i\Delta r_{core}$, $r_{i+1} = (i+1)\Delta r_{core}$, and $r_{i-1} = (i-1)\Delta r_{core}$, yielding the familiar form:

$$\frac{dC_i}{dt} = \frac{\alpha_{core}}{i\Delta r_{core}^2}\left((i+1)C_{i+1} - 2iC_i + (i-1)C_{i-1}\right), \quad i = 1, 2, \ldots, I_{core} - 1 \tag{S4}$$

Repeating the derivation of Equation (S3) for $R_{core}/R_{shell} < r < 1$ gives

$$\frac{dC_j}{dt} = \frac{\alpha_{shell}}{r_j \Delta r_{shell}}\left(C_{j+1} - C_{j-1}\right) + \frac{\alpha_{shell}}{\Delta r_{shell}^2}\left(C_{j+1} - 2C_j + C_{j-1}\right), \quad j = 1, 2, \ldots, J_{shell} - 1 \tag{S5}$$

or

$$\frac{dC_j}{dt} = \frac{\alpha_{shell}}{\Delta r_{shell}^2}\left(\frac{r_{j+1}}{r_j}C_{j+1} - 2\frac{r_j}{r_j}C_j + \frac{r_{j-1}}{r_j}C_{j-1}\right), \quad j = 1, 2, \ldots, J_{shell} - 1 \tag{S6}$$

Unlike in the core, the coordinates in the shell are $r_j = R_{core}/R_{shell} + j\Delta r_{shell}$, $r_{j+1} = R_{core}/R_{shell} + (j+1)\Delta r_{shell}$, and $r_{j-1} = R_{core}/R_{shell} + (j-1)\Delta r_{shell}$. Thus, Equation (S5) is adopted in Equation (13) for the interior points $j = 1, 2, \ldots, J_{shell} - 1$ in the shell layer with $r_j = I_{core}\Delta r_{core} + j\Delta r_{shell}$. The compact forms in Equations (S3) and (S6) are used in the derivation of the interface equations.

*S1.2. Interface*

As the interface condition defined in Equations (4) and (5) allows for two different values at the interface, we need to devise schemes for calculating both of these values. We adopt the fictitious node method [1, 2], which was used to derive the scheme for the boundary condition at the center of the sphere (Equation (12) for $i = 0$). Figure S1 illustrates the discrete points near the interface, with an additional grid point at $r_{i+1}$ that is considered to extend the core past the interface by $\Delta r_{core}$ into the shell and another at $r_{j-1}$ that extends the shell beyond the interface by $-\Delta_{shell}$ into the core. Then, the interior point schemes from Equations (S3) and (S6) can be used. However, the values of the fictitious points need to be determined. We use the interface condition to determine equations for these points.

We apply the normalized radial position and insert $\alpha$ from Equation (10) into Equation (4) to yield

$$\alpha_{core}\frac{\partial C_{core}(R_{core}/R_{shell}, t)}{\partial r} = \alpha_{shell}\frac{\partial C_{shell}(R_{core}/R_{shell}, t)}{\partial r} = F \tag{S7}$$

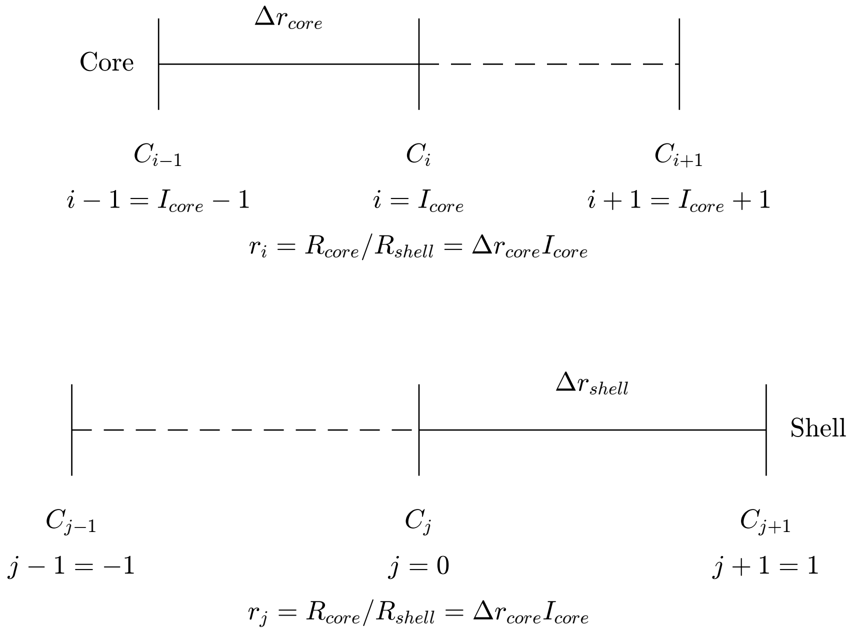


Figure S1: Fictitious node method applied to the interface between two concentric layers of a bi-layered sphere. $C_i$ and $C_j$ denote the concentrations at the interface in the core and shell layers, respectively. The fictitious points are $C_{i+1}$ and $C_{j-1}$. $\Delta r_{core}$ and $\Delta r_{shell}$ are not shown to scale; the key point is that they do not need to be equal.

where $F$ is the continuous flux at the interface $r = R_{core}/R_{shell}$. Using the centered difference approximation around $i = I_{core}$ and $j = 0$ for the core and the shell, respectively, at the interface (Figure S1) gives

$$\alpha_{core}\left(\frac{C_{i+1}-C_{i-1}}{2\Delta r_{core}}\right) = \alpha_{shell}\left(\frac{C_{j+1}-C_{j-1}}{2\Delta r_{shell}}\right) = F \tag{S8}$$

Equation (S8) can be rearranged to solve for the fictitious points $C_{i+1}$ and $C_{j-1}$ as functions of $F$:

$$C_{i+1} = C_{i-1} + \frac{2F\Delta r_{core}}{\alpha_{core}} \tag{S9}$$

$$C_{j-1} = C_{j+1} - \frac{2F\Delta r_{shell}}{\alpha_{shell}} \tag{S10}$$

Inserting Equations (S9) and (S10) into Equations (S3) and (S6), respectively, at the interface ($i = I_{core}$ and $j = 0$) gives

$$\frac{dC_i}{dt} = \frac{\alpha_{core}}{\Delta r_{core}^2}\left(\frac{r_{i+1}}{r_i}\left(C_{i-1} + \frac{2F\Delta r_{core}}{\alpha_{core}}\right) - 2\frac{r_i}{r_i}C_i + \frac{r_{i-1}}{r_i}C_{i-1}\right), \quad i = I_{core} \tag{S11}$$

and

$$\frac{dC_j}{dt} = \frac{\alpha_{shell}}{\Delta r_{shell}^2}\left(\frac{r_{j+1}}{r_j}C_{j+1} - 2\frac{r_j}{r_j}C_j + \frac{r_{j-1}}{r_j}\left(C_{j+1} - \frac{2F\Delta r_{shell}}{\alpha_{shell}}\right)\right), \quad j = 0 \tag{S12}$$

These equations simplify to

$$\frac{dC_i}{dt} = \frac{\alpha_{core}}{\Delta r_{core}^2}\left(2\left(C_{i-1} - C_i\right) + \frac{r_{i+1}}{r_i}\left(\frac{2F\Delta r_{core}}{\alpha_{core}}\right)\right) \tag{S13}$$

and

$$\frac{dC_j}{dt} = \frac{\alpha_{shell}}{\Delta r_{shell}^2}\left(2\left(C_{j+1} - C_j\right) - \frac{r_{j-1}}{r_j}\left(\frac{2F\Delta r_{shell}}{\alpha_{shell}}\right)\right) \tag{S14}$$

Rearranging to solve for $F$ in both equations and equating them gives

$$F = \frac{r_i\alpha_{core}}{2r_{i+1}\Delta r_{core}}\left(\frac{dC_i}{dt}\frac{\Delta r_{core}^2}{\alpha_{core}} - 2\left(C_{i-1} - C_i\right)\right) = \frac{r_j\alpha_{shell}}{2r_{j-1}\Delta r_{shell}}\left(2\left(C_{j+1} - C_j\right) - \frac{dC_j}{dt}\frac{\Delta r_{shell}^2}{\alpha_{shell}}\right) \tag{S15}$$

Now that $F$ has been eliminated, the partition coefficient from Equation (5) can be applied, i.e., $C_i = \kappa C_j$ at the interface. Also, applying derivatives with respect to time at the interface, we have that

$$\frac{dC_i}{dt} = \kappa\frac{dC_j}{dt}, \quad i = I_{core}, j = 0 \tag{S16}$$

Equation (S16) is adopted as the scheme at the interface in the core side (Equation (12) at $i = I_{core}$).

Inserting Equation (S16) into Equation (S15) yields

$$\frac{r_i\alpha_{core}}{2r_{i+1}\Delta r_{core}}\left(\kappa\frac{dC_j}{dt}\frac{\Delta r_{core}^2}{\alpha_{core}} - 2\left(C_{i-1} - \kappa C_j\right)\right) = \frac{r_j\alpha_{shell}}{2r_{j-1}\Delta r_{shell}}\left(2\left(C_{j+1} - C_j\right) - \frac{dC_j}{dt}\frac{\Delta r_{shell}^2}{\alpha_{shell}}\right) \tag{S17}$$

Inserting $\gamma$ defined as

$$\gamma = \frac{\alpha_{shell}\,\Delta r_{core}}{\alpha_{core}\,\Delta r_{shell}} \tag{S18}$$

and moving all the $r$ values to the right-hand side gives

$$\kappa\frac{dC_j}{dt}\frac{\Delta r_{core}^2}{\alpha_{core}} - 2\left(C_{i-1} - \kappa C_j\right) = \gamma\frac{r_j r_{i+1}}{r_{j-1} r_i}\left(2\left(C_{j+1} - C_j\right) - \frac{dC_j}{dt}\frac{\Delta r_{shell}^2}{\alpha_{shell}}\right) \tag{S19}$$

Letting $\Gamma$ be defined as

$$\Gamma = \gamma\frac{r_{i+1}}{r_{j-1}}, \tag{S20}$$

remembering that $r_i = r_j$ at the interface, and rearranging to group like terms gives

$$\frac{dC_j}{dt}\left(\kappa\frac{\Delta r_{core}^2}{\alpha_{core}} + \Gamma\frac{\Delta r_{shell}^2}{\alpha_{shell}}\right) = 2\Gamma C_{j+1} - 2\left(\Gamma + \kappa\right)C_j + 2C_{i-1} \tag{S21}$$

Solving for the derivative at the interface yields

$$\frac{dC_j}{dt} = \frac{2\Gamma C_{j+1} - 2\left(\Gamma + \kappa\right)C_j + 2C_{i-1}}{\kappa\frac{\Delta r_{core}^2}{\alpha_{core}} + \Gamma\frac{\Delta r_{shell}^2}{\alpha_{shell}}}, \quad j = 0 \text{ and } i = I_{core} \tag{S22}$$

which is solved simultaneously with Equation (S16) to find the concentrations at the interface from the shell side and the core side, respectively. Equation (S22) is adopted as the scheme at the interface in the shell side (Equation (13) at $j = 0$).

#### *S1.3. Interface initial condition*

For the initial condition at the interface, we approximate Equation (S7) with first-order accurate forward and backward differences around $i = I_{core}$ and $j = 0$ at the interface (Figure S1) to yield only 1 unknown $C_j(0)$ instead of the three unknowns that would result from the fictitious node method applied for $t > 0$ in Section S1.2:

$$\alpha_{core}\left(\frac{C_i(0) - C_{i-1}(0)}{\Delta r_{core}}\right) = \alpha_{shell}\left(\frac{C_{j+1}(0) - C_j(0)}{\Delta r_{shell}}\right) \tag{S23}$$

Substituting $\gamma$ from Equation (S18) and rearranging yields

$$C_i(0) - C_{i-1}(0) = \gamma\left(C_{j+1}(0) - C_j(0)\right) \tag{S24}$$

Replacing $C_i(0)$ with Equation (5) gives

$$\kappa C_j(0) - C_{i-1}(0) = \gamma\left(C_{j+1}(0) - C_j(0)\right) \tag{S25}$$

Rearranging for $C_j(0)$ gives the initial condition at the interface in the shell:

$$C_j(0) = \frac{\gamma C_{j+1}(0) + C_{i-1}(0)}{\kappa + \gamma}, \quad i = I_{core} \text{ and } j = 0 \tag{S26}$$

Then, the initial condition at the interface in the core is

$$C_i(0) = \kappa C_j(0) = \kappa\frac{\gamma C_{j+1}(0) + C_{i-1}(0)}{\kappa + \gamma}, \quad i = I_{core} \text{ and } j = 0 \tag{S27}$$

*S1.4. Grid spacing*

We have non-uniform grid spacing in the region near the interface between the core and the shell. For grid points $r_i$ for $i \leq I_{core} - 1$ and $r_j$ for $j \geq 1$, the grid spacing is uniform at $\Delta r_{core}$ and $\Delta r_{shell}$, respectively. This means that the equations for the interior points only depend on $\Delta r_{core}$ (Equation (S5)) or $\Delta r_{shell}$ (Equation (S4)), not their combination. The non-uniform spacing in the zone adjacent to the interface (Figure S1) leads to Equation (S22) and thus Equation (S16) having dependence on $\Delta r_{core}$ and $\Delta r_{shell}$ at $r_i = r_j = R_{core}/R_{shell}$ with $i = I_{core}$ and $j = 0$.

We selected $\Delta r_{core}$ and $\Delta r_{shell}$ based on a relationship shown in the literature [3, 4] to maintain 2nd order accuracy of the centered-difference approximation applied to spatial second derivatives when using a non-uniform grid:

$$\Delta r_{shell} - \Delta r_{core} = O(\Delta r_{core}^2) \tag{S28}$$

where $O$ is the big-O notation denoting being of the order of a quantity, instead of being precisely equivalent to the quantity.

In our MATLAB code, we first selected the desired number of spatial intervals in the domain as $M_{desired} = 300$, which would result in grid spacing of $1/M_{desired}$ if the entire bi-layered sphere with normalized outer radius of 1 were uniformly spaced. Then, we set the number of spatial intervals in the core $M_{core}$ based on a proportional split along the total radius:

$$M_{core} = \texttt{floor}\left(M_{desired}\frac{R_{core}}{R_{shell}}\right) \tag{S29}$$

where the `floor` function in MATLAB rounds down to the nearest integer. Then, the grid spacing in the core is

$$\Delta r_{core} = \frac{R_{core}}{R_{shell}}\frac{1}{M_{core}} \tag{S30}$$

Next, Equation (S28) is applied and solved for an approximate grid spacing in the shell layer:

$$\Delta r_{shell,approx} = \Delta r_{core}^2 + \Delta r_{core} \tag{S31}$$

Equation (S31) gives an approximate value because we have to use an integer number of intervals in the shell $M_{shell}$ to have uniform spacing throughout the layer. We compute $M_{shell}$ by

$$M_{shell} = \texttt{ceil}\left(\frac{1}{\Delta r_{shell,approx}}\frac{R_{shell} - R_{core}}{R_{shell}}\right) \tag{S32}$$

where the `ceil` function in MATLAB rounds up to the nearest integer. Finally, the actual $\Delta r_{shell}$ value used is calculated as

$$\Delta r_{shell} = \frac{1}{M_{shell}}\frac{R_{shell} - R_{core}}{R_{shell}} \tag{S33}$$

When $R_{core} = 5.10$ µm of chitosan and $\Delta R = 1.25$ µm of PCL, the resulting values are $M_{core} = 240$, $M_{core} = 59$, $\Delta r_{core} = 3.346456692913386 \times 10^{-3}$, and $\Delta r_{shell} = 3.336447350860694 \times 10^{-3}$, which are close but not equivalent to $1/M_{desired} = 1/300 = 3.333333333333334 \times 10^{-3}$. The grid spacing values are updated for any user-defined inputs of $R_{core}$ and $R_{shell}$ based on the procedure detailed in this section. If the user desires to ensure $M_{core} + M_{core} = M_{desired}$, then the `floor` function in Equation (S29) could be switched to `ceil`. This switch did not change the results presented in the main text in any detectable way.

### S2. Model verification

The finite element model and the finite difference model were verified against each other for the arbitrary case where $D_{core} = 1 \times 10^{-14}$ cm$^2$/s, $D_{shell} = 1 \times 10^{-13}$ cm$^2$/s, $B = 5\%$, $\kappa = 1$, and $\gamma = 10.03$. To this end, we compared drug concentrations at four distinct locations within the core-shell microspheres. Results showed that the drug concentration profiles in both MATLAB and COMSOL overlapped at the center of the microsphere ($r = 0$, Figure S2a), at half the core radius ($r = R_{core}/2$, Figure S2b), and at halfway through the shell layer ($r = R_{core} + \Delta R/2$, Figure S2d). At initial time points, there was a concentration difference at the interface ($r = R_{core}$, Figure S2c) related to the software implementations of the initial condition at the interface (Equation (S26) = Equation (S27) for $\kappa = 1$ for MATLAB and implicitly defined in COMSOL); however, the concentration profiles quickly converged towards the same values after the algorithms started the solver routines (Figure S2c).

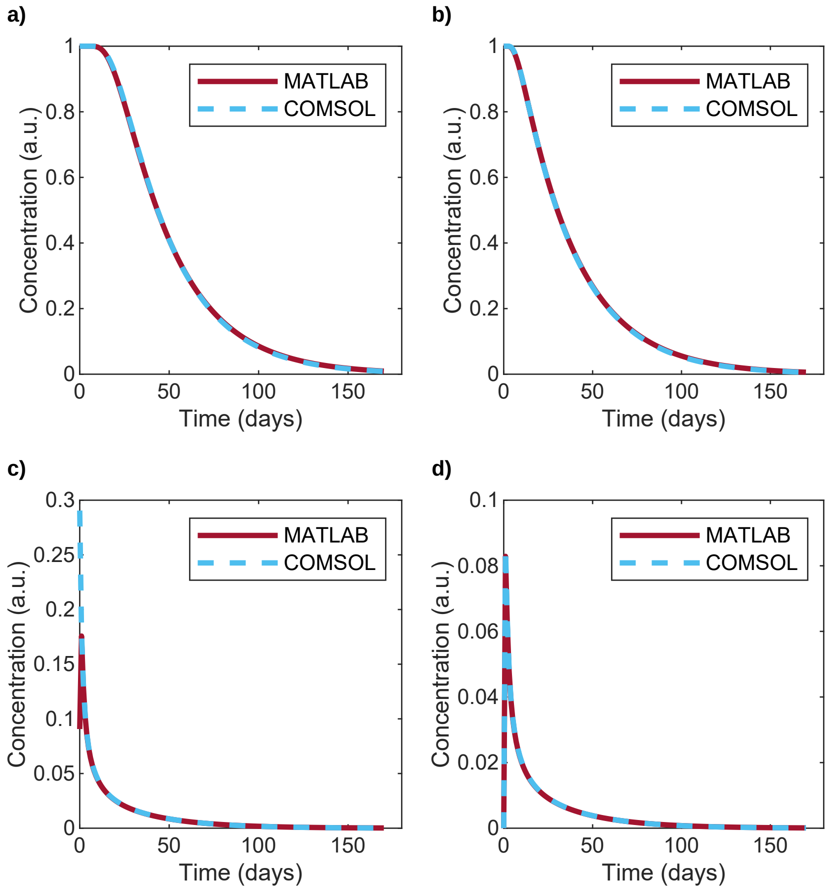


Figure S2: Drug concentration comparison at a) the center of the microsphere ($r = 0$), b) half the core radius ($r = R_{core}/2$), c) the core-shell interface ($r = R_{core}$), and d) half the shell layer thickness ($r = R_{core} + \Delta R/2$). Arbitrary case with $D_{Chi} = 1 \times 10^{-14}$ cm$^2$/s, $D_{PCL} = 1 \times 10^{-13}$ cm$^2$/s, $B = 5\%$, $\kappa = 1$, and $\gamma = 10.03$. $D_{Chi} = D_{core}$: drug diffusion coefficient in the chitosan core. $D_{PCL} = D_{shell}$: drug diffusion coefficient in the polycaprolactone shell. $B$: burst release. $\kappa$: partition coefficient. $\gamma$: ratio of diffusion coefficients and grid spacing in the core and shell layers defined by Equation (S18).

### S3. Preliminary parameter estimation

To avoid converging to a local minimum, we employed a multi-start approach with 100 randomized initial parameter guesses generated using the Latin hypercube sampling method with uniform sampling across the log of each parameter. The preliminary parameter range boundaries were defined based on the following criteria:

- The lower limit for the burst was set to no immediate release ($B = 0$).
- The upper limit for burst was set to the theoretical complete immediate release ($B = 100\%$).
- The lower and upper limits for the diffusion coefficients $D_{Chi}$ and $D_{PCL}$ were set based on feasible values that bounded the experimental data from Jiang et al. [5] between simulated results (Figure S3) considering $B = 0$, $\kappa = 1$, and $D_{Chi} = D_{PCL}$.
- The upper limit for $D_{PCL}$ was set to $5 \times 10^{-11}$ cm$^2$/s to expand the range to allow for faster release through the PCL shell.
- The limits for the partition coefficient were chosen arbitrarily.

The parameters' lower and upper limits for the preliminary parameter estimation are tabulated in Table S1.

Table S1: Limits used in the preliminary parameter estimation.

| **Parameter** | **Lower limit** | **Upper limit** | **Units** |
|---|---|---|---|
| $B$ | 0 | 100 | % |
| $D_{Chi}$ | $1 \times 10^{-15}$ | $1 \times 10^{-13}$ | cm$^2$/s |
| $D_{PCL}$ | $1 \times 10^{-15}$ | $5 \times 10^{-11}$ | cm$^2$/s |
| $\kappa$ | 0.1 | 10 | - |

$B$: burst release. $D_{Chi} = D_{core}$: drug diffusion coefficient in the chitosan core. $D_{PCL} = D_{shell}$: drug diffusion coefficient in the PCL shell. $\kappa$: partition coefficient.

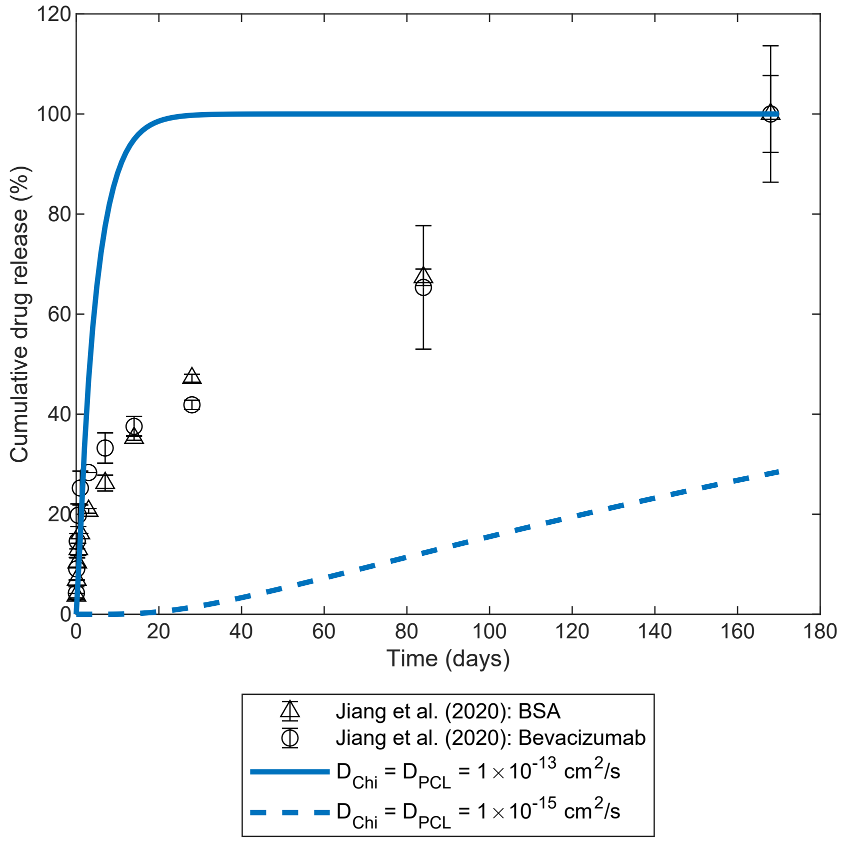


Figure S3: Cumulative drug release profiles determining feasible lower and upper limits for $D_{Chi}$ and $D_{PCL}$ when considering core-shell microsphere with drug loaded in the core only. Experimental data from Jiang et al. [5]. $D_{Chi} = D_{core}$: drug diffusion coefficient in the chitosan core. $D_{PCL} = D_{shell}$: drug diffusion coefficient in the polycaprolactone shell.

After completing the 100 multi-start parameter estimations, we evaluated the resulting error values (Figure S4a,b for BSA and Figure S5a,b for bevacizumab). We arbitrarily set a cut-off value at a 5% error increase relative to the minimum error obtained (Table S2). We considered all the simulations with errors below this threshold as part of the acceptable parameters set used to estimate average parameters. Furthermore, the parameters from the simulation with the lowest error value were considered the best model parameters from the preliminary parameter estimation phase. Parameters from 53 out of 100 MATLAB simulations and 95 out of 100 COMSOL simulations were used to calculate the average parameters of the model for BSA. For bevacizumab, the final parameters from 64 out of 100 MATLAB simulations and 95 out of 100 COMSOL simulations were used to obtain the parameters for the average model in each software. From the parameter results obtained within the error threshold, a new set of values for the upper limits of the parameters was defined to reduce the search space.

For MATLAB, the average and best (minimum error over all the multi-start parameter estimations) model predictions overlapped, and the COMSOL model had some variation between the best and average model predictions for BSA and overlapped for bevacizumab (Figure S4c,d for BSA and Figure S5c,d for bevacizumab).

The last experimental data point shows a standard error exceeding 100% release, which may impair the models' ability to predict the later time points accurately. This experimental cumulative release above 100% could be attributed to several factors. One potential reason is the sensitivity of the instrument used for measuring the drug release at later time points. Another possibility is an inaccurate calculation of the initial amount of the drug in the microspheres due to the way the loading efficiency was measured.

Table S2 contains the values of the parameters for the average and best models in both software programs for both drugs. Comparing the best models in the software programs and the average models to the best models shows that the parameters obtained for burst release and BSA diffusion coefficient in chitosan were similar; however, a greater difference was observed between the results for BSA diffusion coefficient in PCL and the partition coefficient. Even so, the error values of the best results in both software programs were similar.

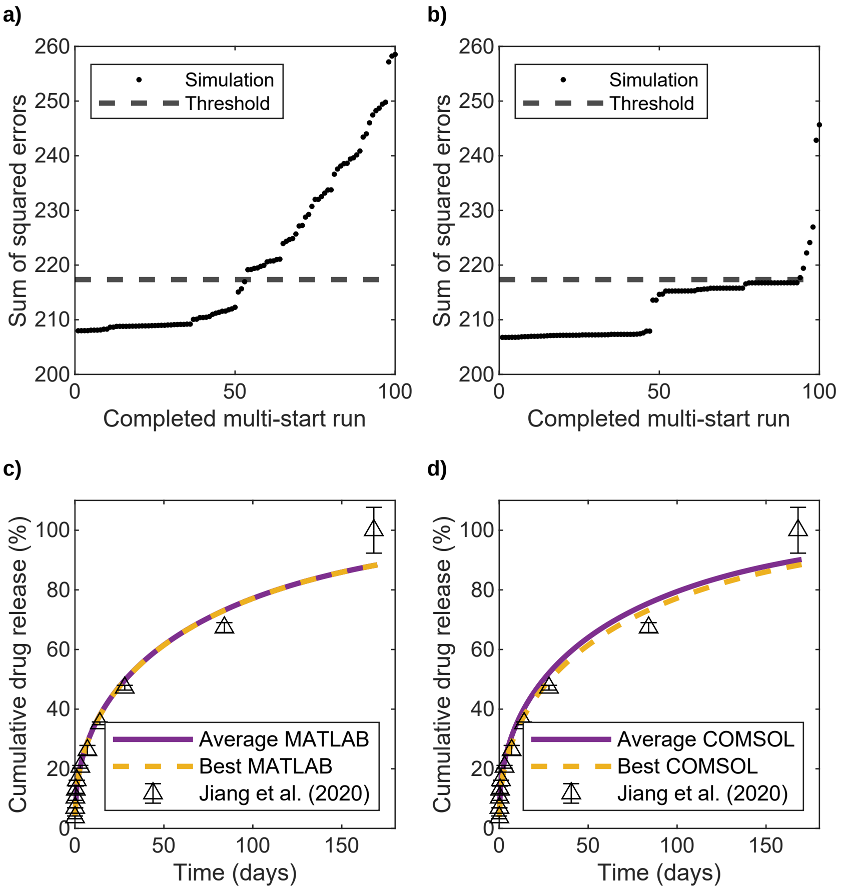


Figure S4: Error values and cumulative drug release profiles for BSA release after the preliminary parameter estimation. a) Error values sorted in increasing order from MATLAB. b) Error values sorted in increasing order from COMSOL. c) Average and best MATLAB models compared to experimental data. d) Average and best COMSOL models compared to experimental data. Experimental data from Jiang et al. [5].

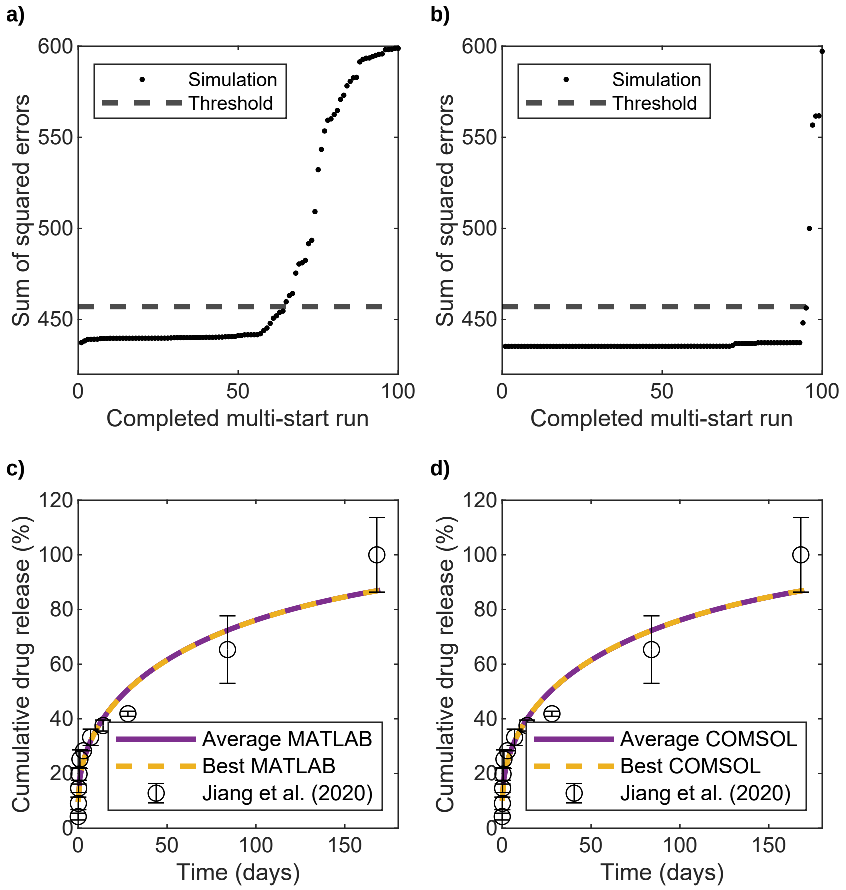


Figure S5: Error values and cumulative drug release profiles for bevacizumab release after preliminary parameter estimation. a) Error values in MATLAB. b) Error values in COMSOL. c) Average and best MATLAB models compared to experimental data. d) Average and best COMSOL models compared to experimental data. Experimental data from Jiang et al. [5].

Table S2: Preliminary parameter estimation results: estimated parameters for fitting the model to data for drug-loaded chitosan-PCL core-shell microspheres from Jiang et al. [5]. The average model was obtained by averaging the parameters obtained from all the optimization runs that achieved an error within 5% of the minimum error after 100 multi-start attempts. The best model corresponds to the parameter set with the lowest error value.

| **BSA in MATLAB** | | | | | |
|---|---|---|---|---|---|
| **Average model** | | | **Best model** | | |
| **Parameter** | **Value** | **Units** | **Parameter** | **Value** | **Units** |
| $B$ | 4.30 | % | $B$ | 4.01 | % |
| $D_{Chi}$ | $2.91\times10^{-15}$ | $cm^2/s$ | $D_{Chi}$ | $2.90\times10^{-15}$ | $cm^2/s$ |
| $D_{PCL}$ | $4.17\times10^{-12}$ | $cm^2/s$ | $D_{PCL}$ | $2.96\times10^{-11}$ | $cm^2/s$ |
| $\kappa$ | 7.53 | - | $\kappa$ | 8.69 | - |
| Error value | 210.11 | - | Error value | 207.98 | - |
| **BSA in COMSOL** | | | | | |
| **Average model** | | | **Best model** | | |
| **Parameter** | **Value** | **Units** | **Parameter** | **Value** | **Units** |
| $B$ | 6.12 | % | $B$ | 3.81 | % |
| $D_{Chi}$ | $3.15\times10^{-15}$ | $cm^2/s$ | $D_{Chi}$ | $2.91\times10^{-15}$ | $cm^2/s$ |
| $D_{PCL}$ | $1.85\times10^{-12}$ | $cm^2/s$ | $D_{PCL}$ | $3.13\times10^{-12}$ | $cm^2/s$ |
| $\kappa$ | 7.79 | - | $\kappa$ | 1.38 | - |
| Error value | 211.54 | - | Error value | 206.77 | - |
| **Bevacizumab in MATLAB** | | | | | |
| **Average model** | | | **Best model** | | |
| **Parameter** | **Value** | **Units** | **Parameter** | **Value** | **Units** |
| $B$ | 9.80 | % | $B$ | 10.12 | % |
| $D_{Chi}$ | $2.62\times10^{-15}$ | $cm^2/s$ | $D_{Chi}$ | $2.54\times10^{-15}$ | $cm^2/s$ |
| $D_{PCL}$ | $6.14\times10^{-12}$ | $cm^2/s$ | $D_{PCL}$ | $5.64\times10^{-12}$ | $cm^2/s$ |
| $\kappa$ | 6.50 | - | $\kappa$ | 1.97 | - |
| Error value | 441.12 | - | Error value | 437.27 | - |
| **Bevacizumab in COMSOL** | | | | | |
| **Average model** | | | **Best model** | | |
| **Parameter** | **Value** | **Units** | **Parameter** | **Value** | **Units** |
| $B$ | 10.05 | % | $B$ | 10.01 | % |
| $D_{Chi}$ | $2.58\times10^{-15}$ | $cm^2/s$ | $D_{Chi}$ | $2.58\times10^{-15}$ | $cm^2/s$ |
| $D_{PCL}$ | $1.07\times10^{-12}$ | $cm^2/s$ | $D_{PCL}$ | $1.07\times10^{-12}$ | $cm^2/s$ |
| $\kappa$ | 2.47 | - | $\kappa$ | 1.05 | - |
| Error value | 436.06 | - | Error value | 435.28 | - |

$B$: burst release. $D_{Chi} = D_{core}$: drug diffusion coefficient in the chitosan core. $D_{PCL} = D_{shell}$: drug diffusion coefficient in the polycaprolactone shell. $\kappa$: partition coefficient.

### S4. Spatial concentration distributions

Figure S6 shows the normalized drug concentration profiles across the normalized radius of the core-shell microsphere at different time points for the three distinct regimes: a) core-limited diffusion ($D_{Chi} << D_{PCL}$), b) balanced diffusion ($D_{Chi} = D_{PCL}$), and c) shell-limited diffusion ($D_{Chi} >> D_{PCL}$). In the core limited regime (Figure S6a), the steepest slopes appear in the core, with a relatively low and constant concentration in the shell due to the fast outward diffusion. In the case of equal diffusion coefficients (Figure S6b), the concentration gradients are smooth and symmetric across the core and shell. In the shell-limited regime (Figure S6c), a constant concentration is observed in the core with a sharp drop in the shell due to the slow outward diffusion. The relative diffusivities shape the spatial and temporal distribution of drugs within DDSs and can be used to inform design strategies that tune drug release. It should be noted that these plots are consistent with those from [6] for normalized radial concentration profiles in bi-layered spheres.

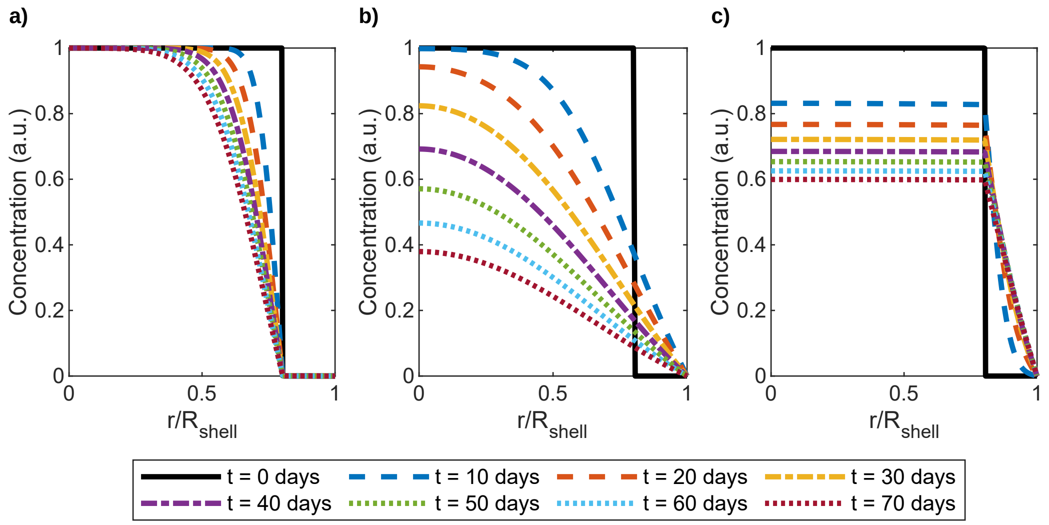


Figure S6: Drug concentration profiles for different time points as a function of normalized position. For each case, $B = 10$ and $\kappa = 1$. First column: $D_{Chi} << D_{PCL}$. Second column: $D_{Chi} = D_{PCL}$. Third column: $D_{Chi} >> D_{PCL}$. a) $D_{Chi} = 1 \times 10^{-15}$ cm$^2$/s and $D_{PCL} = 1000 \times D_{Chi} = 1 \times 10^{-12}$ cm$^2$/s, b) $D_{Chi} = 1 \times 10^{-14}$ cm$^2$/s and $D_{PCL} = D_{Chi}$, and c) $D_{Chi} = 1 \times 10^{-12}$ cm$^2$/s and $D_{PCL} = 0.001 \times D_{Chi} = 1 \times 10^{-15}$ cm$^2$/s. $B$: burst release. $\kappa$: partition coefficient. $D_{Chi} = D_{core}$: drug diffusion coefficient in the chitosan core. $D_{PCL} = D_{shell}$: drug diffusion coefficient in the polycaprolactone shell.

### S5. Sensitivity analysis

Figure S7 shows the results for the MOAT screenings performed in COMSOL. The calculated MOAT mean measures the effect of each parameter on the cumulative drug release, and the MOAT standard deviation estimates the nonlinear effect of each parameter and the interaction effect of each parameter with other parameters. In the case where $D_{Chi} << D_{PCL}$ (Figure S7a), the drug diffusion coefficient in the chitosan core was the most influential parameter, followed by burst release. In contrast, the drug diffusion in the PCL shell and the partition coefficient had a negligible impact on the cumulative drug release at 28 days. When $D_{Chi} = D_{PCL}$ (Figure S7b), the sensitivities changed, and the drug diffusion coefficient in PCL became the most sensitive parameter, followed by the partition coefficient and the drug diffusion coefficient in chitosan. In this case, burst release had minimal impact on the cumulative drug release at 28 days. For the case where $D_{Chi} >> D_{PCL}$ (Figure S7c), the burst release was the dominant parameter, followed by the drug diffusion coefficient in PCL and the drug partition coefficient. The drug diffusion coefficient in chitosan had a negligible effect on the cumulative drug release at 28 days. In all three cases, none of the parameters had significant interactions with the other parameters as the MOAT standard deviations ($y$-axis values) were small (Figure S7).

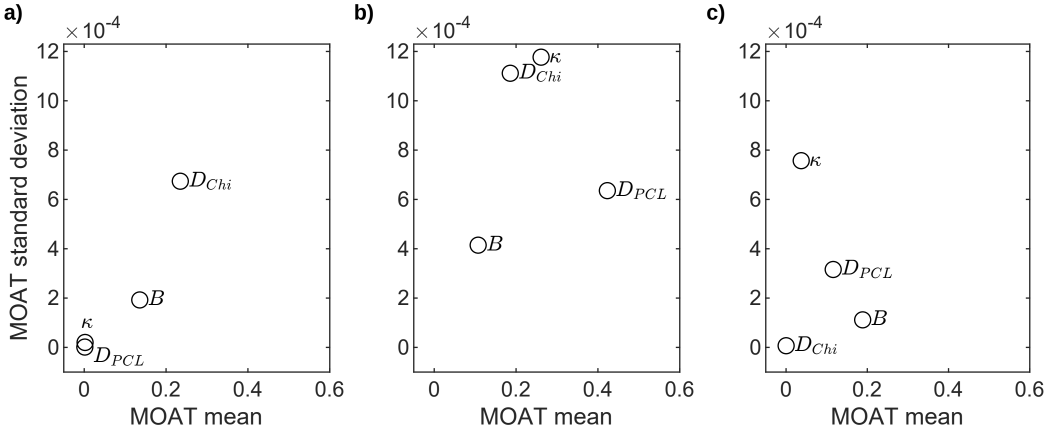


Figure S7: Morris one-at-a-time (MOAT) method for assessing the effects of the different parameters on cumulative drug release at 28 days using the finite element solution in COMSOL. Each parameter was considered to have a normal distribution with a 1% standard deviation. For each case, $B = 10$ and $\kappa = 1$. First column: $D_{Chi} << D_{PCL}$. Second column: $D_{Chi} = D_{PCL}$. Third column: $D_{Chi} >> D_{PCL}$. a) $D_{Chi} = 1 \times 10^{-15}$ cm$^2$/s and $D_{PCL} = 1000 \times D_{Chi} = 1 \times 10^{-12}$ cm$^2$/s, b) $D_{Chi} = 1 \times 10^{-14}$ cm$^2$/s and $D_{PCL} = D_{Chi}$, and c) $D_{Chi} = 1 \times 10^{-12}$ cm$^2$/s and $D_{PCL} = 0.001 \times D_{Chi} = 1 \times 10^{-15}$ cm$^2$/s. $B$: burst release. $\kappa$: partition coefficient. $D_{Chi} = D_{core}$: drug diffusion coefficient in the chitosan core. $D_{PCL} = D_{shell}$: drug diffusion coefficient in the polycaprolactone shell. MOAT mean is an estimate of the overall effect of the parameter on cumulative drug release. MOAT standard deviation is a measure of nonlinear effects of the parameter and its interaction with other parameters.

### S6. Predictive capabilities extended for twice the duration and half the release rate threshold

In Section 3.3 we simulated the model for 180 days and showed the drug release rate threshold of 2 μg/day and the window of ± 10% around that threshold for Figures 5–7. To consider the case of drug release rate threshold of 1 μg/day, we also needed to extend the simulation duration, so we doubled the duration to 360 days. The following figures are analogous to Figures 5–7 but with results until 360 days and drug release rate threshold of 1 μg/day and the window of ± 10% around that threshold.

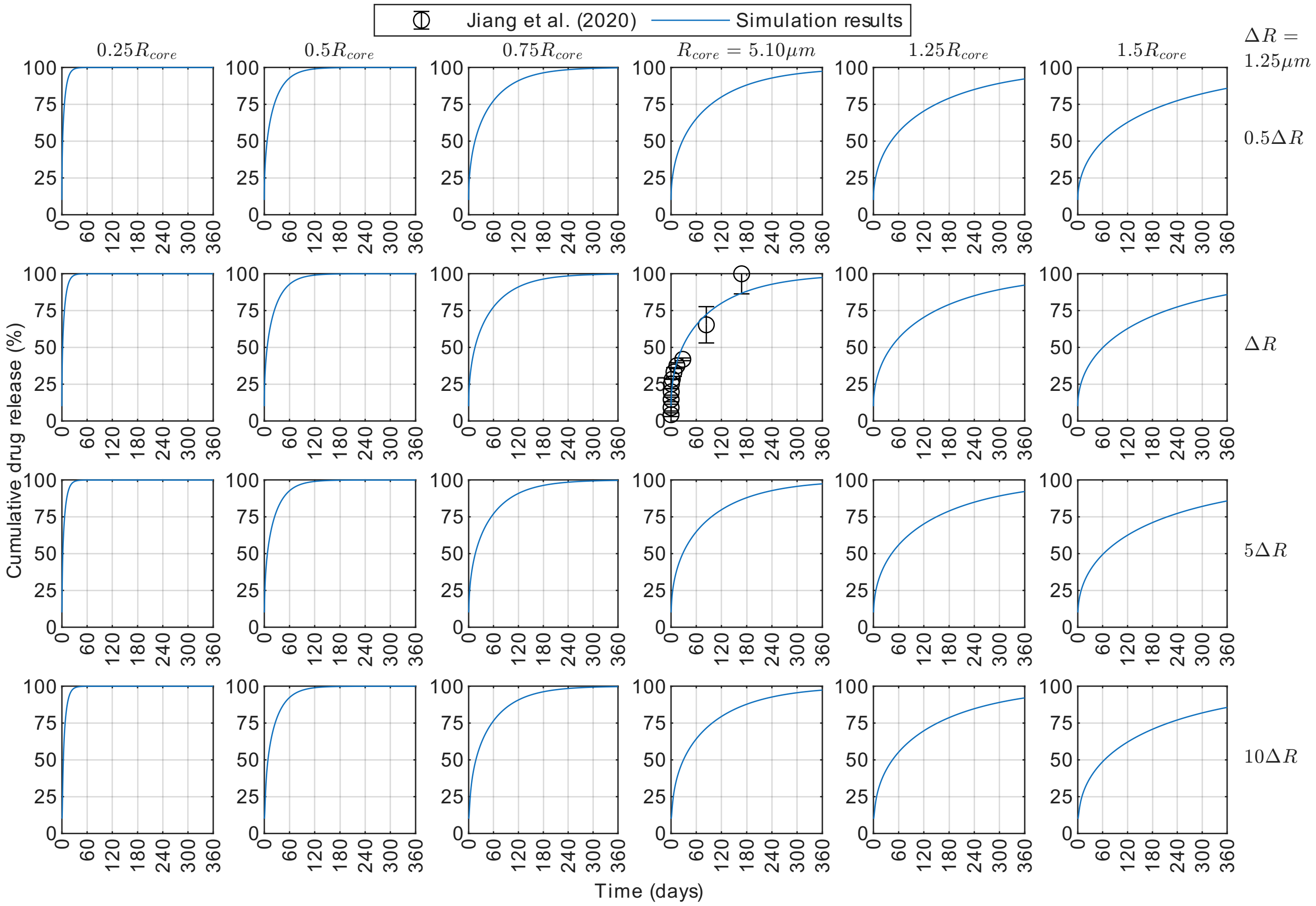


Figure S8: Cumulative drug release profiles until 360 days for bevacizumab released from core-shell microspheres with drug loaded in the core only. Results were obtained in MATLAB with $D_{Chi} = 2.6 \times 10^{-15}$ cm$^2$/s, $D_{PCL} = 2.6 \times 10^{-12}$ cm$^2$/s, $B = 10\%$, and $\kappa = 1$ (average model parameters from Table 2). Experimental data for $R_{core} = 5.10$ μm and $\Delta R = R_{shell} - R_{core} = 1.25$ μm are from Jiang et al. [5]. Each panel shows profiles for different chitosan-PCL configurations where the core radius and shell thickness are varied by multipliers to the baseline dimensions $R_{core}$ and $\Delta R$ considered from Jiang et al. [5]. The core radii are labeled on the columns, and the shell thicknesses are labeled on the rows. $D_{Chi} = D_{core}$: drug diffusion coefficient in the chitosan core. $D_{PCL} = D_{shell}$: drug diffusion coefficient in the polycaprolactone shell. $B$: burst release. $\kappa$: partition coefficient.

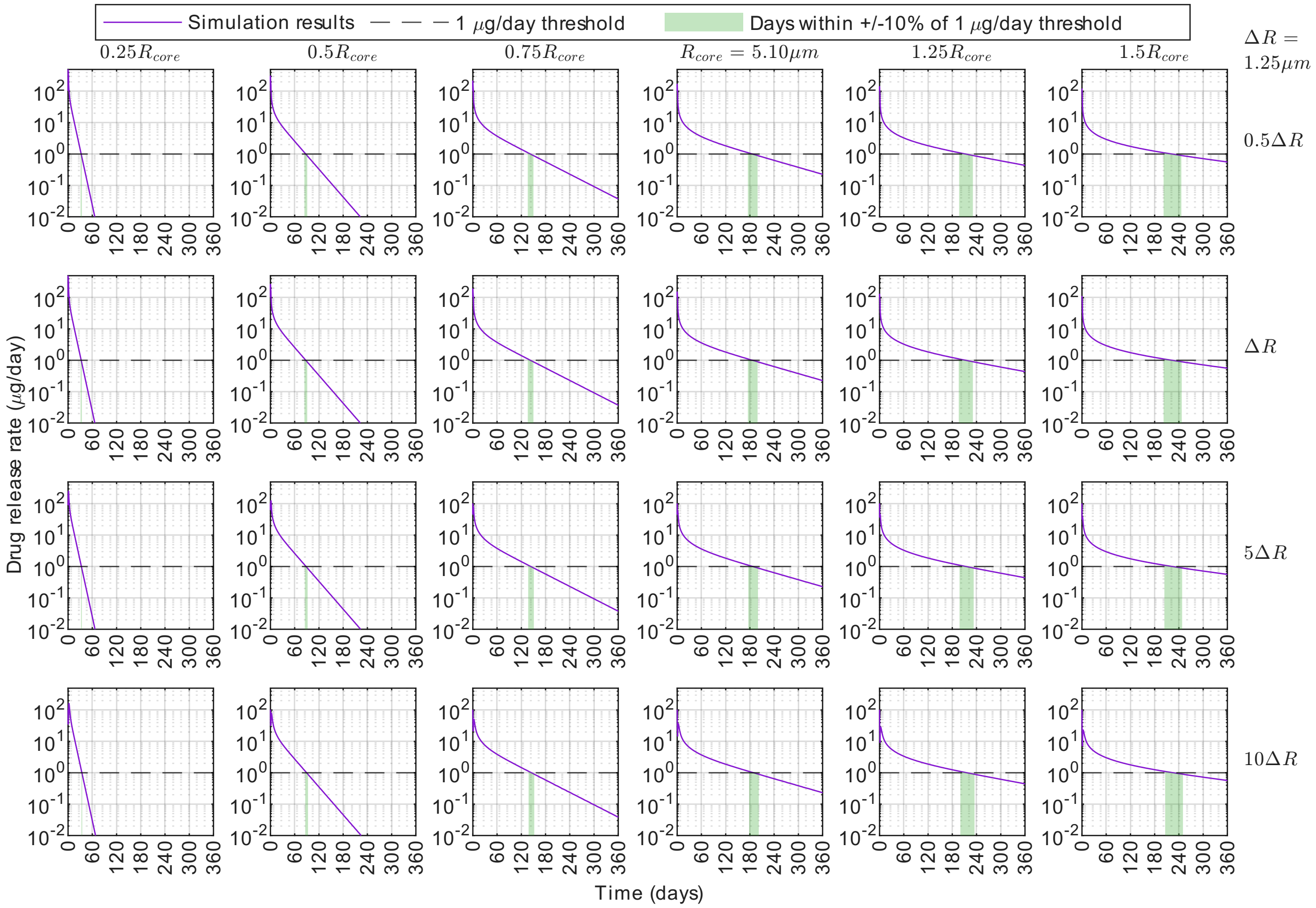


Figure S9: Drug release rate profiles until 360 days for bevacizumab released from core-shell microspheres with drug loaded in the core only. Results were obtained in MATLAB with $D_{Chi} = 2.6 \times 10^{-15}$ cm$^2$/s, $D_{PCL} = 2.6 \times 10^{-12}$ cm$^2$/s, $B = 10\%$, and $\kappa = 1$ (average model parameters from Table 2). Each panel shows profiles for different chitosan-PCL configurations where the core radius and shell thickness are varied by multipliers to the baseline dimensions $R_{core} = 5.10$ µm and $\Delta R = R_{shell} - R_{core} = 1.25$ µm considered from Jiang et al. [5]. The core radii are labeled on the columns, and the shell thicknesses are labeled on the rows. On each panel, the purple curve is the simulation results, the dashed line shows the 1 µg/day drug release rate threshold, and the shaded green region highlights the days within ± 10% of the 1 µg/day drug release rate threshold. $D_{Chi} = D_{core}$: drug diffusion coefficient in the chitosan core. $D_{PCL} = D_{shell}$: drug diffusion coefficient in the polycaprolactone shell. $B$: burst release. $\kappa$: partition coefficient.

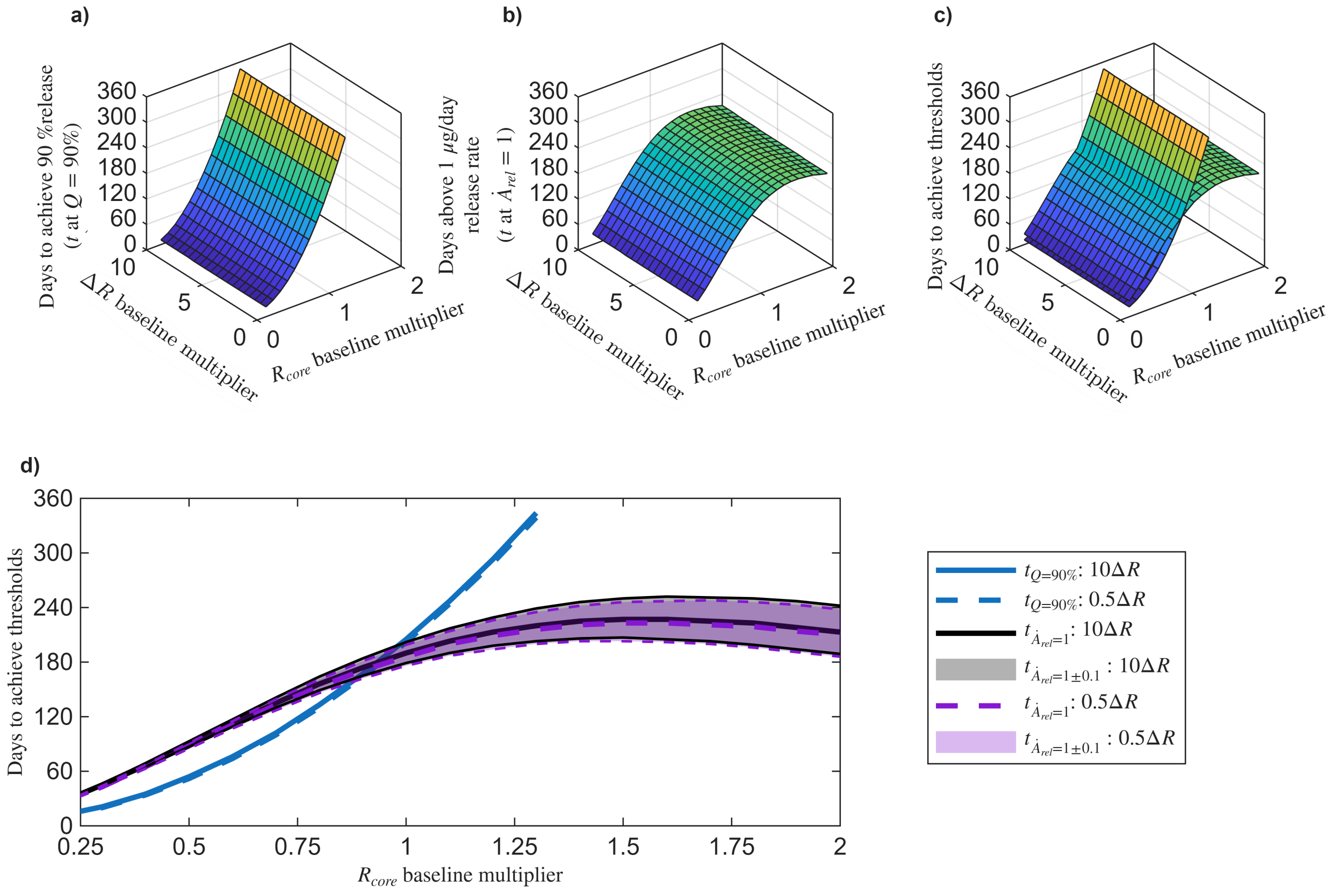


Figure S10: Time in days to reach cumulative release ($Q$) and release rate ($\dot{A}_{rel}$) thresholds for different chitosan-PCL configurations (including those from Figures S8 and S9) where the core radius and shell thickness are varied by multipliers to the baseline dimensions $R_{core} = 5.10$ μm and $\Delta R = R_{shell} - R_{core} = 1.25$ μm considered from Jiang et al. [5]. a) Time in days to reach cumulative release threshold of $Q = 90\%$ as a function of $R_{core}$ and $\Delta R$ baseline multipliers. b) Time in days to reach release rate threshold of $\dot{A}_{rel} = 1$ μg/day as a function of $R_{core}$ and $\Delta R$ baseline multipliers. c) 3D view of the surfaces from panels a) and b) combined. d) 2D view of panel c) in the projection onto the time vs. $R_{core}$ baseline multiplier axes. $\Delta R$ baseline multipliers of 0.5 and 10 are shown in panel d) along with shaded regions denoting the intervals where the release rates are within ± 10% of the threshold. Results for all panels were obtained in MATLAB with $D_{Chi} = 2.6 \times 10^{-15}$ cm$^2$/s, $D_{PCL} = 2.6 \times 10^{-12}$ cm$^2$/s, $B = 10\%$, and $\kappa = 1$ (average model parameters from Table 2). $R_{core} = 5.10$ μm and $\Delta R = R_{shell} - R_{core} = 1.25$ μm are from Jiang et al. [5]. $D_{Chi} = D_{core}$: drug diffusion coefficient in the chitosan core. $D_{PCL} = D_{shell}$: drug diffusion coefficient in the polycaprolactone shell. $B$: burst release. $\kappa$: partition coefficient.
